# Personalized whole-brain Ising models with heterogeneous nodes capture differences among brain regions

**DOI:** 10.1101/2025.06.09.658769

**Authors:** Adam Craig, Sida Chen, Qianyuan Tang, Changsong Zhou

## Abstract

Multiple lines of research have studied how complex brain dynamics emerge from underlying connectivity by using Ising models as simplified neural mass models. However, limitations on parameter estimation have prevented their use with individual, high-resolution human neuroimaging data. Furthermore, most studies focus only on connectivity, ignoring node heterogeneity, even though real brain regions have different structural and dynamical properties.

Here we present an improved approach to fitting Ising models to 360-region functional MRI data: derivation of an initial guess model from group data, optimization of simulation temperature, and two stages of Boltzmann learning, first with group data, then with individual data. Our implementation uses GPU acceleration to mitigate the high computational cost of this approach. We then analyze how data binarization threshold affects goodness-of-fit, the role of the external field in model behavior, consistency among models fitted to different scans of the same individual, and correlations between model parameters and features from structural MRI, including measures of myelination and cortical folding.

We find that binarizing fMRI data at higher thresholds decreases correlation between model and data functional connectivity but increases the heterogeneity of node external fields and their correlations with structural features. A choice of threshold that achieves both goodness-of-fit and intrinsic heterogeneity of regions results in a model that better reflects the reality of the brain as a network of intrinsically heterogeneous nodes. By enabling personalized, biophysically interpretable modeling of structure-function mapping across the whole brain, this approach can aid understanding of individual differences in brain network organization and bridge the gap between the network-focused methodology of connectomics and the region-focused paradigm typical of translational research.

## 1. Introduction

### 1.1. The heterogeneous brain

Understanding how individual differences in brain structure shape differences in brain dynamics remains a central goal of modern neuroscience. The study of connectomics represents one prominent paradigm, in which researchers have made significant progress in linking individual differences in structural connectivity (SC), functional connectivity (FC), and cognitive abilities (Zimmermann et al., 2018; Wang et al., 2019b; Liégeois et al., 2020). However, the brain is not just a network of interchangeable nodes. Large-scale brain network analyses have revealed that microstructural features, such as myelination or cortical granularity, can significantly modulate local excitation-inhibition balance and structure-function coupling across brain regions (Glasser et al., 2014; Fotiadis et al., 2023; Fotiadis et al., 2024). Heterogeneity of local excitability can in turn influence larger-scale brain dynamics (Abeywardena, 2024). Importantly, individual differences in the distribution of structural features across the brain can also predict differences in cognitive traits (Kristanto et al., 2023), suggesting that brain regions influence behavior through both their locations in the larger network and through their intrinsic properties. However, the high dimensionality of MRI data and the complexity of integrating single-subject functional and structural data make translating from correlations to mechanistic insights challenging (Bijsterbosch, et al., 2020; Venkadesh & Van Horn, 2021; Sultana et al., 2024).

### 1.2. The need for explainable, data-driven models

To meet this need, network-based approaches have shifted from static structure-function correlations to dynamical models with varying degrees of personalization (Lynn & Bassett, 2019; Guo et al., 2025). Such models could enable simulations of how the brain will respond to stimulation of different brain regions, leading to therapies tailored to the needs of each patient (Lynn & Bassett, 2019; Guo et al., 2025).

One extreme example of the detail now possible in dynamical models of the human brain is the Digital Twin Brain Project, which simulated 86 billion leaky integrate-and-fire neurons with 47.8 trillion synapses informed by multiple kinds of *in vivo* human data (Lu et al., 2024). However, it also illustrates the challenges of such a model, such as the risk of overfitting a high-dimensional model to limited data and the impracticality of comparing simulations of multiple individuals. Even before reaching this scale, the modeling community realized the need for approximations at macroscopic scales and developed multiple ways of modeling collective neural behavior, including neural mass models (Breakspear, 2017, Bansal et al., 2018). Both early neural mass models of animal nervous systems (Zhao et al., 2010) and newer models based on human brain data have suggested that regional heterogeneity and SC together shape patterns of FC (Demirtaş et al., 2019, Wang et al., 2019a, Deco et al., 2021).

However, these also face tradeoffs, as more complex models suffer from lower reliability despite greater individual specificity (Domhof et al., 2022). One key problem is that functional MRI (fMRI) provides an accurate view of spatial correlations but not temporal correlations in the underlying brain activity (Tu et al., 2024). This suggests that using a simpler model that captures spatial patterns of integration and segregation but is robust against spurious temporal fluctuations could provide a better bridge between brain structure and brain function.

### 1.3. Ising models as neural mass models

Over the past decade, the Ising model has proven capable of modeling many kinds of systems, because, when fitted to data, such models can capture the mean activity of and pairwise correlations between variables (Nguyen & Zecchina, 2017). After fitting, their simplicity makes their behavior explainable and their parameters amenable to a wide variety of analyses (Nguyen & Zecchina, 2017). Most relevantly, a growing number of researchers have used all-to-all connected (generalized) Ising models as simplified neural mass models. When diffusion-MRI-derived SC serves as the network of couplings between regions, simulated FC correlates with fMRI-derived empirical FC, albeit modestly (correlation < 50%) (Das et al., 2014; Abeyasinghe et al., 2018). Despite the limitations of this approach, researchers have used it to model information transfer between nodes (Marinazzo et al., 2014), disorders of consciousness (Abeyasinghe et al., 2020), and the effects of anesthesia (Stramaglia et al., 2017), audio stimulation (Kandeepan et al., 2020), and sedatives (Kandeepan et al., 2020).

Other studies have achieved better correlations between model and data FC by fitting the model parameters directly to fMRI data, or, less commonly, to intracranial electroencephalogram data (Ashourvan et al., 2021). Notable downstream analyses of fMRI-fitted Ising models include clustering model states by the energy barriers between them (Ezaki & Watanabe, 2017), identifying important subnetworks through parameter sensitivity (Ponce-Alvarez et al., 2022), and calculating the correlation between the coupling parameters and SC (Ruffini et al., 2023). However, these examples pooled data from multiple individuals due to the data-intensive nature of the pseudolikelihood maximization fitting method they employed. This limitation to population-level models prevents the study of individual differences.

This inability to fit Ising models to individual data represents a major shortcoming compared to maximum likelihood gradient ascent, which other studies have used to fit models to single-subject fMRI data (Watanabe et al., 2013; Theis et al., 2024; Theis et al., 2025). These models have shown consistent and diagnostically meaningful individual differences in energy distribution among states (Theis et al., 2024; Theis et al., 2025). However, due to the need to compute the energies of all possible states, the authors only modeled small networks of fewer than 10 regions. Bayesian approximations allow fitting of larger 15-region models to individual data and show similar goodness of fit (Khanra et al., 2024). Parameters from individual models fitted with Bayesian approximations correlated significantly (around 50%-60%) with cognitive scores (Kang et al., 2021) or attention deficit and hyperactivity disorder-associated traits (Jeong et al., 2021). This apparent significance of individual differences in Ising model parameters suggested that being able to use a more fine-grained parcellation to identify areas of interest will improve the utility of Ising models in future practical applications. Specifically, many of the most promising targets for treating psychiatric disorders with transcranial magnetic stimulation are in higher-order areas, such as the prefrontal cortex and show high inter-patient variability in SC, making precise, personalized targeting valuable (Cash & Zalesky, 2024).

Choice of binarization threshold for the fMRI time series is an additional, insufficiently explored consideration. Recent work showed that threshold choice can reshape the inferred energy landscape even in 10-node models (Theis et al., 2025), suggesting that it could have an even greater impact at whole-brain scales. Works that have addressed alternate choices of threshold have mainly considered them in terms of how they affected goodness of fit (Watanabe et al., 2013; Fortel et al., 2022; Theis et al., 2025), not how thresholding influenced the external fields, which are potentially important as the only distinguishing intrinsic features of nodes.

More generally, the role of the external field in the Ising model has received relatively little attention. Studies that use the SC for the coupling matrix typically drop the external field from the model entirely (Das et al., 2014; Marinazzo et al., 2014; Stramaglia et al., 2017; Abeyasinghe et al., 2018; Abeyasinghe et al., 2020; Fortel et al., 2022), though (Kandeepan et al., 2020) uses it to represent an external stimulus. No prior work involving Ising models fitted to fMRI data has specifically examined the importance of heterogeneity of external field to goodness of fit (Watanabe et al., 2013; Ezaki et al., 2017; Jeong et al., 2021; Kang et al., 2021; Ponce-Alvarez, 2022; Ruffini et al., 2023; Khanra et al., 2024; Theis et al., 2024; Theis et al., 2025). The studies that considered structure-function relationships only compared couplings and SC (Watanabe et al., 2013; Fortel et al., 2022; Ruffini et al., 2023). Of the two studies that use sparse canonical correlation analysis to compare individualized Ising model parameters to cognitive or behavioral scores, (Kang et al., 2021) uses the external field and couplings together, while (Jeong et al., 2021) uses only the couplings. The heterogeneity of nodes in the fitted Ising model represents an overlooked opportunity to connect structural features to local excitability and local excitability to whole-brain-scale dynamics.

### 1.4. Personalized whole-brain Ising models with heterogeneous nodes

Here we present a GPU-accelerated, four-stage workflow that fits fully connected 360-region Ising models to individual HCP resting-state fMRI data, which we then apply to an analysis of the influence of binarization threshold on goodness of fit, node heterogeneity, and correlations between external field and brain structural features. Whereas we use correlation between fMRI data-derived FC and model-derived FC to verify goodness of fit between the model and the data to which we fitted it, we demonstrate the biological relevance of the model by comparing the model couplings to DTI data-derived SC and the model external fields to structural MRI features. This comparison of both region-wise and pair-wise model parameters to structural features reflects the reality that the regions of the brain have both their own distinct features and relationships to other regions. To achieve this, we search for a fMRI data binarization threshold that balances both goodness of fit to FC and heterogeneity model external fields.

Our approach builds on Boltzmann learning with Metropolis Monte Carlo sampling, which others have used for fitting Ising models to spiking neuron recordings (Roudi et al., 2009). While computationally intensive, it offers a high degree of reliability when given sufficient sampling simulation length and parameter update steps (Roudi et al., 2009). We improved the reliability of the method and decreased the number of fitting steps needed by starting from a data-derived initialization and optimizing simulation temperature. We also made use of GPU acceleration to generate hundreds of personalized models in parallel.

To quantify how binarization threshold influences goodness of fit and the role of heterogeneous external fields, we first compared different thresholds applied to the same pooled group data. As in (Watanabe et al., 2013; Fortel et al., 2022; Theis et al., 2025), we found that fitting was fastest and most reliable when we binarized data at the mean, at which threshold we also observed a correlation between model coupling and SC close to that reported by (Ruffini et al., 2023). However, we found that fitting succeeded at thresholds up to 1 SD above the mean and that this higher threshold was necessary to elicit meaningful node heterogeneity that correlated with region-to-region differences in myelination, curvature, and other cortical features measured with structural MRI. We then used this threshold to fit single-subject models and found that individual differences in external field contributed to capturing individual differences in FC. Finally, we quantified the local structure-function coupling of each region as the predictability of individual differences in Ising model external field from individual differences in structural features.

To the best of our knowledge, this is the first work to demonstrate fitting of 360-region whole-brain Ising models to single-subject data or to use Ising model external field parameters to study the relationship between local excitability and cortical structural features. In summary, our results provide a principled route toward structure-informed, subject-specific Ising models that provide insights into how local microstructure and global network topology jointly sculpt brain functional connectivity, an essential step toward mechanistic biomarkers and precision neuro-stimulation planning.

## 2. Methods

### 2.1. Human brain MRI data

This study is based on the publicly available dataset of the WU-Minn Young Adult Human Connectome Project (HCP), including different brain imaging modalities covering resting-state fMRI, DTI, and structural MRI (Van Essen et al., 2013). The HCP project had been approved by the local Institutional Review Board at Washington University in St. Louis.

The data to which we fit the Ising models is resting-state fMRI data of healthy young adults from the HCP S1200 data release (Elam et al., 2021). We selected 837 individuals who each had a complete set of imaging data, including four fMRI time series, each 1200 time points sampled 0.72 seconds apart (14.4 minutes per scan), a DTI scan, a T1-weighted scan, and a T2-weighted scan. See (Van Essen et al., 2017) for more information about data collection procedures.

We preprocessed the fMRI time series in the manner used in (Wang et al., 2021). This pipeline started with the HCP minimal preprocessing pipeline (Glasser et al., 2013), which performed basic cleaning and alignment. These steps reflect common practice and provide the bare minimal steps needed to compare data across subjects (Glasser et al., 2013). We then assigned voxels to regions in the 360-region parcellation from (Glasser et al., 2016) to produce a 360×1200 time series for each scan. This Atlas divides the brain into regions that are relatively homogenous in both their FC and structural properties (Glasser et al., 2016), which is useful for our goal of comparing Ising model parameters derived from fMRI data to structural MRI features and DTI-derived SC data. Finally, we band-pass filtered each region time series to the infra-slow band (0.01-0.1 Hz), the frequency range in which the blood-oxygenation-level-dependent (BOLD) signal components correlate most strongly with changes in infra-slow electrophysiological oscillations and behavior (Palva & Palva, 2012). For fitting results with the same HCP data parcellated with the 117-region Automatic Anatomical Labeling (AAL) Atlas Version 1 (Tzourio-Mazoyer et al., 2002), see Supplementary Figure S1.

A key stage of creating the fitting targets was binarization of the data, which required selection of a threshold. As in (Theis et al., 2025), we defined the threshold in terms of standard deviations (SD) above the mean of the local region time series and mapped values below the threshold to −1 and values above it to +1. To create an individual fitting target, we concatenated the four binarized time series of individual scans into a single 360 × 4800 sequence and took the means 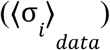 and uncentered covariances 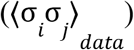. To create a group target, we randomly selected a training set of 670 subjects and took the means and uncentered covariances of their concatenated binarized time series data. We set aside the remaining 167 individuals as the testing set to test whether using the group Ising model as the initial guess for the individualized Ising model fitting would lead to differences in goodness of fit between individuals in and out of the training set. We created group model fitting targets by binarizing the z-scored and pooled fMRI data at 31 binarization thresholds evenly spaced between 0 and 3 SD above the mean, inclusive. Because fitting a separate ensemble of 837 single-subject models at every threshold was computationally impractical, we selected two thresholds, the endpoints of the range of thresholds at which we could reliably fit group Ising models, 0 and 1.

To derive a SC matrix from each DTI scan, we used the second-order integration over fiber orientation distributions ProbtrackX probabilistic tract-tracing method from (Behrens et al., 2004; Behrens et al., 2007) as implemented in the FSL v5.0.9 software (https://fsl.fmrib.ox.ac.uk/fsl/docs/diffusion/probtrackx.html/). We used the same default parameters as in (Wang et al., 2021). Briefly, for each voxel, we sampled 5000 streamlines that started from that voxel and propagated with a step length of 0.55 mm and terminated when they reached the pial surface, encountered an angle of 0.2 degrees between one step and the next, or extended to a maximum tract length of 2000 steps (Wang et al., 2021). We then computed the probability that a streamline starting in one of the 360 regions would pass through each other, producing a 360×360 matrix of probabilities (Wang et al., 2021). We then took the mean of the matrix and its transpose to make the final SC matrix symmetric and set its diagonal to 0 to remove self-connectivity (Wang et al., 2021). In practice, because both the SC and Ising model coupling matrices are symmetric with zeros along the diagonal, we only computed correlations between corresponding elements above the diagonals.

Each region has four structural features derived from T1- and T2-weighted structural MRI data: thickness, myelination, curvature, and sulcus depth. The myelination estimate is the ratio of T1 to T2 values (T1w/T2w ratio) at each voxel, so that the value for a region is the mean over its voxels as in (Kristanto et al., 2023). The remaining features (thickness, curvature, and sulcus depth) are morphometric measurements of the cortical surface rather than of individual voxels that come from the final phase of the FreeSurfer recon-all method (Fischel, 2012) and are available in GIfTI shape files distributed with the HCP S1200 data (Glasser et al., 2013). For these surface mesh properties, we took the mean value over all applicable vertices covering a given region of the parcellation as done in (Kristanto et al., 2023). The cortical thickness is the distance between the pial surface and the gray-white matter border (Kristanto et al., 2023). The remaining two measures reflect folding of the pial surface. The curvature is the bending of each vertex, and the sulcus depth is the distance between vertices at the midpoint of each sulcus and the midpoint of the nearest gyrus (Kristanto et al., 2023).

While many studies have used the T1w/T2w ratio as an estimate of myelination, recent validation studies have suggested that it lacks specificity and has limited agreement with some other measures (Arshad, 2017; Sandrone, 2023). A wide variety of alternative methods for measuring myelination exist, but all have potential sources of error, suggesting that comparing different metrics can provide a more robust characterization (Lazari, 2021). In Supplementary Section S14, we compare parameter correlation results for the T1w/T2w ratio and the combined four T1- and T2-weighted structural MRI feature vector to individual and combined features from the AMICO pipeline (Daducci, 2015), including both standard DTI features and features from the NODDI 2-shell model (Zhang, 2012). Myelin-associated features obtained include the common DTI-derived feature fractional anisotropy (Lazari, 2021) and the neurite density index (Jespersen, 2010).

### 2.2. Ising model simulation and fitting

The generalized Ising model maintained the same formulation used in previous fMRI studies that used them as simplified neural mass models (Ezaki & Watanabe, 2017; Ruffini et al., 2023; Ponce-Alvarez et al., 2022; Theis et al., 2024; Khanra et al., 2024; Theis et al., 2025): Each node *i* corresponded to a brain region and had an external field parameter, *h*_*i*_, which influenced its probability of being in the down-regulated state (σ_*i*_ =− 1) or up-regulated state (σ_*i*_ =+ 1). Each pair of regions (*i, j*) had a coupling parameter, *J*_*ij*_, which influenced the probability of coactivation of two regions and was symmetric (*J*_*ij*_ = *J*_*ji*_) with no self-coupling (*J*_*ii*_ = 0). The energy of a state 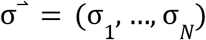 of an *N*-node system, 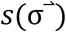, and the partition function *Z* taken over all 2^*N*^ possible states determined the probability 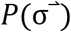:

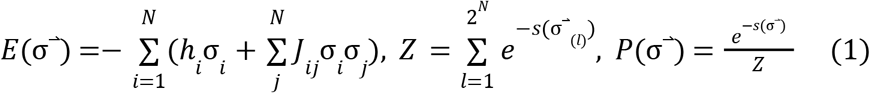

The most direct method of fitting an Ising model to a sampling of states derived from data is likelihood maximization, but this requires *O*(2^*N*^) time per optimization step, as one must calculate the probabilities of all possible states (Ezaki & Watanabe, 2017). The most common alternative is pseudolikelihood maximization, which takes *O*(*TN*^2^) time per optimization step, where *T* is the number of observed states in the data, but this method only approximates true likelihood maximization for large *T* (Ezaki & Watanabe, 2017). Consequently, the studies that use it aggregate data from many individuals, as seen in (Ezaki & Watanabe, 2017; Ruffini et al., 2023; Ponce-Alvarez et al., 2022). For examples of results of applying the pseudo-likelihood maximization method from (Ezaki & Watanabe, 2017) to binarized group-level HCP data, see Supplementary Figure S2.

We instead fitted models using the version of Boltzmann learning described in (Roudi et al., 2009). In this method, parameter updates are proportional to the discrepancies between the means 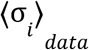 and uncentered covariances 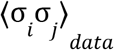 from the target data and those of the model, 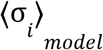 and 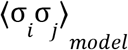 as sampled with Metropolis Monte Carlo simulation (Roudi et al., 2009). The updates to the model parameters are the differences between the first and second order mean fields of the time series generated by the model and those computed from the data, scaled by an arbitrary learning rate ηϵ[0, 1] (Roudi et al., 2009):

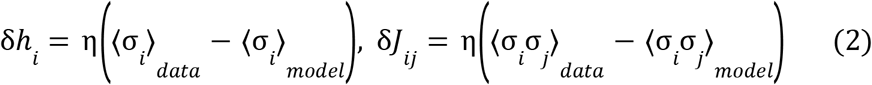

Metropolis simulation was the most computationally costly part of this fitting method, so we used two strategies to make our implementation more efficient. First, we used a sequential update scheme, meaning that we iterated over all the nodes in a fixed order, giving each node state a chance to update, before recording the system state as a new sample. As shown in (Potter & Swendsen, 2013), sequential update maintains total balance while also being the computationally simplest update scheme. In their tests, this allowed it to achieve the fastest convergence to the true probability distribution of states in terms of real-world computation time (Potter & Swendsen, 2013). Second, we used the PyTorch (v1.13.1) framework to run multiple simulations in parallel on a graphics processing unit (GPU). While the Metropolis algorithm requires that we update node states serially, it is straight-forward to run multiple independent simulations of equal-sized systems in parallel using vector arithmetic operations on multi-dimensional arrays.

In addition to simple improvements in computational efficiency, we improved the performance of the Boltzmann learning method by using the mean field values from the data as parameters in an initial guess model and using this initial guess to optimize the simulation temperature. In early tests of individual models, we started from a randomly generated model but found that fitting progressed slowly if at all. By contrast, our initial guess model, after temperature optimization, had a correlation between model and binarized data FC of up to 0.6, which made it possible for the subsequent fitting to converge more quickly. This optimal temperature depended on the choice of binarization threshold and was the main benefit of starting with the initial guess model (See Supplementary Figure S3).

While (Roudi et al., 2009) provides several methods for finding approximate fits to data that one can further improve through Boltzmann learning, we found that the even simpler approach of using the means and uncentered covariances derived from the data provided adequate performance (See Figure 3a):

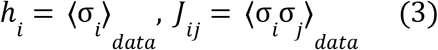

Intuitively, such a model should behave at least qualitatively similarly to the fitted model. Larger positive values of *h*_*i*_ should increase the probability that region *i* will remain in the excited state, increasing 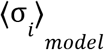, while negative values of *h*_*i*_ should decrease the probability, decreasing 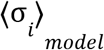. Similarly, larger positive values of *J*_*ij*_ should increase the uncentered covariance between regions *i* and *j*, 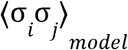, while larger negative values should make them anti-correlated, making 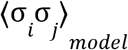, negative. However, this model does not fully reproduce 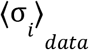 and 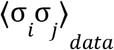 when we sample its states with the Metropolis algorithm, meaning that *δh*_*i*_ and *δJ*_*ij*_ in Equation 2 are initially nonzero and that we can further improve it through Boltzmann learning. While this approach to initialization worked for both group and individual data, we found that using the group model as the initial guess for the individual models lead to faster convergence and better worst-case fits (Supplementary Figure S4), so we used this group-to-individual approach for the individual models presented in subsequent results.

However, in practice, we found that the FC from the Metropolis simulation would only correlate with that of the data when we simulated near the critical inverse temperature (*β*). Researchers have long known that simulating at the critical temperature, also known as the Curie temperature, leads to optimal sampling of the model states (Bian et al., 2010). For all choices of binarization threshold, we found that most regions in the binarized data were positively correlated, making most initial *J*_*ij*_ > 0. Consequently, as *β* increased (temperature decreased) from 0 to some finite freezing point, the model passed from the fully random phase, through a critical phase with frequent phase transitions but with a clear preference for states that align strongly coupled regions, to a frozen phase in one highly aligned configuration.

Our tests showed that the initial guess model only had an FC correlated with that of the data in a narrow window around the critical *β* value (See Figure 3a). To optimize the simulation temperature, we performed a 1-dimensional grid-based search, iteratively homing in on the temperature that produced the lowest root mean squared error (RMSE) between the centered covariances of the initial guess model and the target data. We used the covariance RMSE because, even though it is harder to interpret when comparing goodness of fit of different fitted models, it gives numerically valid results even for frozen or near-frozen phases. By contrast, FC correlation can be undefined when regions have 0 variance (no flips). We first searched over the range [0, 1] for each initial guess group model, sampling 101 evenly spaced values per iteration. In all cases, the temperature converged within 10 iterations, making this step brief compared to the tens of thousands of simulations used for Boltzmann learning. In general, the optimal temperature coincided with a flip acceptance rate close to 0.5, so the flip acceptance rate could serve as a proxy for covariance RMSE that required less computation and had a smaller memory footprint, as it only required tracking of a single value per region instead of a matrix of pairwise covariances. For a comparison, see Supplementary Material section S5.

For the simulate-and-update loop of Boltzmann learning, we used 1200-step simulations, as longer simulations did not improve convergence time or final goodness of fit of the model. However, we did find that longer simulations of 120,000 steps were necessary for simulated FC to converge to a consistent value. To test a model, we computed the Pearson correlation between its FC and the FC of the target binarized data, where we defined the FC as the Pearson correlation between the states of pairs of regions. We ran these longer test simulations once every 1000 parameter updates to identify the point at which fitting ceased to increase the FC correlation. To test the reliability of fitting, we independently fitted 101 replica group models for each fitting target. To ensure that the full set of individual models fit within system memory, we only used 5 replica models for each individual subject fitting target.

We fitted all models for 40,000 parameter updates. We chose a single stopping point for all model fitting runs to allow fair comparison of goodness-of-fit and found that 40,000 updates adequately balanced goodness-of-fit and computation time. The probability of each potential node state update of the Metropolis algorithm depended on the full present state, which made parallelization of a single run of the algorithm impossible. However, we were able to run multiple simulations, whether multiple replicas, binarization thresholds, or subjects, in parallel. Specifically, we used the PyTorch Python library to create an implementation that could run thousands of Ising models (101 replicas of each of 31 thresholds = 3,131 group models; 5 replicas of each of 837 individuals = 4,185 individual models) in parallel on a single nVIDIA Tesla V100 GPU with 32 GB of memory. For both group and individual models, fitting took fewer than 14 days to reach 40,000 parameter updates. To test whether this was adequate to achieve convergence, we compared FC correlations after 40,000 and 100,000 updates and found that, while FC correlations did continue to trend upward, the differences were nearly always small and mainly occurred among the poorly fitting outlier instances. See Supplementary Material Section S6.

While the time required for fitting was substantially higher than that needed for pseudo-likelihood maximization on the group data, which took 24 hours to converge using 80,000 updates, Boltzmann learning required less memory and achieves consistently better fits for binarization thresholds 0 and 1, though not at 2.4 (Supplementary Figure S2). Each run of the pseudolikelihood maximization algorithm required computing the pseudolikelihood using the full time series data, while Boltzmann learning only required storing the mean and uncentered covariance of the data and accumulating the mean and uncentered covariance of the Ising model simulation in-place. The full set of time series binarized at one threshold and concatenated occupied 5.8 GB, while the mean and uncentered covariance only occupied 520 KB. This made it possible to fit more models in parallel on the same GPU.

Both the long fitting time and the substantial memory footprint are consequences of the relatively fine-grained 360-region parcellation along with the all-to-all connectivity of the model. It is possible to substantially reduce the time needed for Boltzmann learning to converge by using a lower-dimensional model. For example, the group model using the 117-region AAL atlas (Supplementary Figure S1) reliably achieved FC correlations over 0.98 at threshold 1 after only 5 days and 20,000 parameter updates. For either Atlas, this reflects the time needed to fit 3,100 models, making the time per model closer to 2.3 minutes per model for 117 regions and 6.5 minutes per model 360 regions. Fitting a single 117-node model takes less than 7 hours, and fitting a single 360-node model takes less than 20 hours on a desktop computer with one NVIDIA GeForce RTX 2070 graphics card. For single-model fitting, any benefit from GPU parallelization is smaller than the overhead. We can roughly halve these times by switching from GPU to CPU. Finally, we translated the same fitting algorithm into a MATLAB script for single-model fitting, which was faster than the Python implementation, requiring only 2.7 hours to run 40,000 parameter updates of a 360-node model or 20 minutes for 40,000 updates of a 117-node model.

To test the importance of region heterogeneity to the behavior of the Ising model, we compared the FC correlation of each model to that of a modified model with homogenized external fields. For the group model test, we used a modified fitting process in which we forced all *h*_*i*_ to the same value by initializing them to a single value 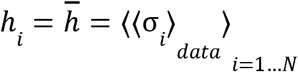 and then updating them with the modified update step 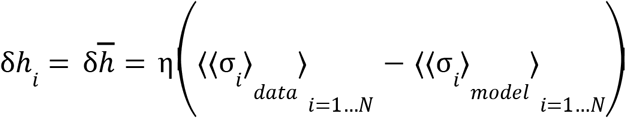, where 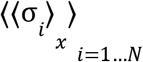 indicates the mean state over regions. To test the importance of individual differences, we instead allowed region-wise differences but modified the individual model fitting to force 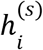 to remain the same across individuals. The modified update step is 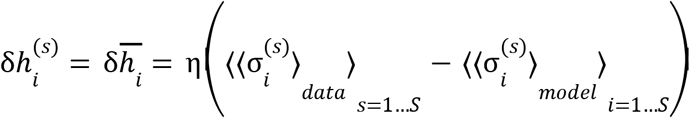, where 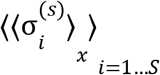 indicates the mean state over individuals. Due to limitations on available computing resources, we were only able to fit the same-h individual models for 35,000 parameter updates, so we compared them to different-h individual models fitted for 35,000 updates.

To test the consistency of Ising models fitted to different fMRI scans of the same individual, we fitted a separate Ising model to each single scan. We then performed a series of comparisons for each pair of scans of the same individual and for each pair of scans of different individuals. For each pair, we computed the region-wise correlation between *h*_*i*_ values, the region-pairwise correlation between *J*_*ij*_ values, the region-pairwise correlation between FC values generated by Metropolis simulation of the model, and the region-pairwise correlation between FC values computed directly from the binarized data. This gave us 4 pairs of distributions to compare. We expected that, in each case, intra-person correlations would be higher than inter-person correlations, so we compared the distributions using 1-sided Mann-Whitney U-tests with Bonferroni-corrected 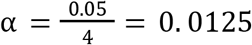. We also tested whether the model and data intra-person FC correlations were different from each other using a 2-sided Wilcoxon signed rank test with α = 0. 05.

### 2.3. Predictability of model parameters from brain structural features

After fitting Ising models to data, we used least squares linear regression models to test for linear relationships between structural features of the brain and Ising model parameters. To avoid biasing results by having multiple replicas of the same model, we took the mean of each parameter over replicas. We considered four sets of linear models:

1. Group model *J*_*ij*_ predicted from the group mean SC, *Ĵ* (*SC*_*ij*_) (1 correlation per model).
2. Group model *h*_*i*_ predicted from the group means of the region features (thickness, myelination, curvature, and sulcus depth), *ĥ* (*t*_*i*_, *m*_*i*_, *c*_*i*_, *d*_*i*_), plus predictions from individual features: *ĥ* (*t*_*i*_), *ĥ* (*m*_*i*_), *ĥ* (*c*_*i*_), and *ĥ* (*d*_*i*_).
3. For each region pair (*i, j*), a separate linear model 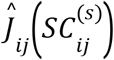 fitted to individual Ising model 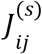 parameters and 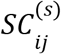 data values of all subjects *s* (360·(360 − 1)/2 = 64, 620 correlations per model, one for each region pair).
4. For each region *i*, a separate linear model 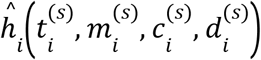 fitted to individual Ising model 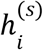 and structural feature values of all subjects *s* plus single-feature models, 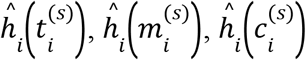, and 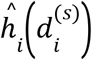 (360 · 5 = 1800 correlations per model, 4 single-predictor regressions and one 4-predictor regression per region).

To demonstrate the value of using the model parameters instead of comparing structural feature data directly to fMRI data, we compared each linear regression model to a similar model predicting a feature derived directly from the binarized fMRI data, either the FC predicted from the SC or the mean state of the binarized region time series from the region features:

1. Group fMRI data *FC*_*ij*_ predicted from the group mean 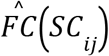 (1 correlation per model).
2. Group fMRI data ⟨σ_*i*_ ⟩ predicted from the group means of the region features (thickness, myelination, curvature, and sulcus depth), 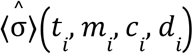, plus predictions from individual features: 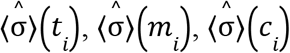, and 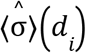.
3. For each region pair (*i, j*), a separate linear model 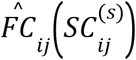 fitted to individual data 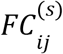 and 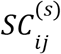 data values of all subjects *s* (360·(360 − 1)/2 = 64, 620 correlations per model, one for each region pair).
4. For each region *i*, a separate linear model 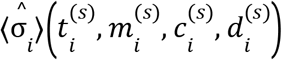 fitted to individual Ising model 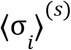 and structural feature values of all subjects *s* plus single-feature models, 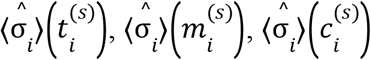, and 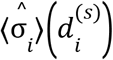 (360 · 5 = 1800 correlations per model, 4 single-predictor regressions and one 4-predictor regression per region).

We then looked for spatial trends in the local standard deviation (SD) of model parameters and the strength of the local individual-level prediction correlations. Hypothesizing that a lack of individual variation in brain structure might explain why some regions had poor predictability of individual model 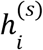 from structure, we computed the correlations between the local SD of each model parameter and the local prediction correlation. For the same reason, we computed correlations between the local SD of SC and the prediction correlation of individual model 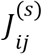. Finally, since several lines of research, including those described in (Fotiadis et al., 2023) and (Paquola et al., 2019), have suggested that regions with greater myelination trade more efficient signal transmission for lower synaptic plasticity or functional variability, we tested whether mean myelination negatively correlated with individual variability (SD) of 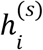.

### 2.4. Testing the significance of correlations and differences

For each correlation, we used a permutation test to test the probability of observing a spurious correlation between two uncorrelated variables with the same distributions. When considering the correlation between a value and a prediction of that value from a least-squares linear regression model, we repeated the linear model fitting for each permutation. In the Results section, we used *p* < 10^−6^ to denote that none of the 10^6^ permuted correlations were stronger than the original.

In cases where we performed a set of similar tests and selected the cases that were significant, we performed a Bonferroni correction by dividing the pre-correction α, 0.05, by the number tests in the set to get a corrected α. For the group model correlations, we initially tested 31 choices of binarization threshold. In most plots, we omitted the upper end of the range, because Ising model fitting failed for some very high thresholds. However, whenever we performed a statistical test for each threshold and selected the significant results, we accounted for the comparison of all 31 thresholds.

For the *J*_*ij*_-*Ĵ*_*ij*_ (*SC*_*ij*_) and 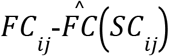 correlations, we further corrected for selecting from these two sets of 31 correlations by using 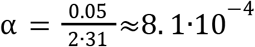. For the group model *h*_*i*_ and group data ⟨σ_*i*_ ⟩ prediction correlations, we considered the four single-feature linear regressions plus the four-feature linear regression, so we used 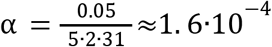. For individual model correlations where we compared prediction correlations of both 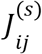 and 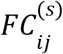 for all 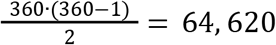 region pairs at thresholds 0 and 1, we used 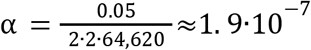. For individual model correlations where we compared predictions of both 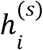 and ⟨σ _*i*_⟩^(*s*)^ at all 360 regions at thresholds 0 and 1, we considered both single-feature and four-feature models, so we used 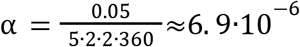.

Finally, we tested the significance of spatial trends in individualized model parameter variability and predictability. We considered correlations between local individual-level correlation strength and local mean and SD of structural features: 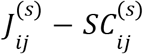 correlation versus mean and SD of 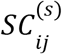 and mean and SD of 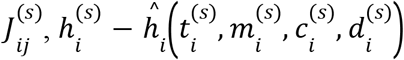 correlation versus mean and SD of each individual feature, 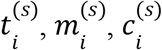, and 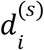, and SD of 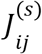 versus the mean and SD of each of the same features. We only consider individual models at binarization threshold 1, so we only needed to correct for 4 + 2·2·5 = 24 correlations and use 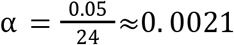.

When repeating these tests with an extended set of features that included six additional brain region features derived from DTI data (Supplementary Section 14), we repeated the permutation tests using new α values to account for the larger number of correlations tested. Since our goal was to test whether the additional DTI features predicted trends in *h*_*i*_ better than the structural MRI features, we did not include correlations with mean binarized state. For the region-wise correlations between group model *h*_*i*_ and mean feature values, we compared the original four T1- and T2-weighted structural MRI features, the six DTI features, and five multiple least-squares regressions with different subsets of features so that 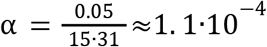. For the extended subject-wise region-local individual model correlations, we considered all ten individual features and three multiple least-squares regressions so that 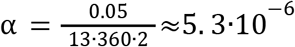. We also performed pairwise correlations between overall distributions of correlations. For threshold 1, we tested whether structural MRI or DTI prediction correlations were greater with a 2-tailed Wilcoxon signed rank test, as we did not expect either to be greater *a priori*. We also tested whether combining both sets of features increased prediction correlations over using just one or the other set using two 1-tailed Wilcoxon tests. For each feature or combination of features, we compared correlations for models with binarization thresholds 0 and 1 using 1-tailed Wilcoxon signed rank tests, as we expected threshold 1 to produce stronger correlations in all cases. Since we perform 16 Wilcoxon tests, we use 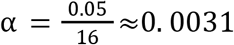.

When checking whether one set of correlations was greater than another, we used 1-sided, 2-sample Wilcoxon signed rank tests. When comparing multiple such tests, we again applied a Bonferroni correction based on the number of tests from which we selected cases with significant differences. When we tested the effect of homogenizing *h*_*i*_ on the behavior of the group models, we considered each binarization threshold, so we used 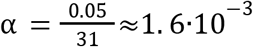. When we tested whether individual models fit individual data better than group models or tested the effect of homogenizing individual model 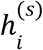 overall subjects *s*, we only considered two binarization thresholds, 0 and 1, so we used 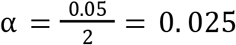. One key difference between the group model tests and the individual model tests is that, whereas, for group model tests, the replicas for the same threshold are the samples, for the individual model tests, we take the mean FC correlation over replicas so that we have one sample per subject.

For comparisons between individual model 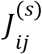 and data 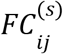, our goal was to test whether one threshold or the other resulted in stronger structure-parameter correlations and whether either threshold lead to 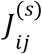 prediction correlations that were stronger than 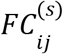 correlations. we considered four pairwise comparisons of sets of correlations: 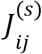 at thresholds 0 and 1, 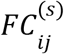 at thresholds 0 and 1, 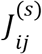 and 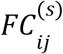 at threshold 0, and 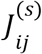 and 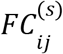 at threshold 1, so we use 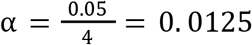. For the individual model 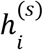 and ⟨σ _*i*_⟩^(*s*)^ prediction correlations, we performed the same 4 pairwise comparisons but repeated them for each single structural feature and for the four-feature vector, so we used 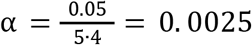.

## 3. Results

### 3.1. Effect of binarization of data on functional connectivity

We first searched for the interval of thresholds over which binarization preserved the overall pattern of FC. Figure 2 shows how binarizing the data affects activity in an example time series (a), alongside covariance (b) and FC (c). Comparing (b) and (c) showed that covariance diminished due to the diminishing variances of the individual region time series, while FC, which was normalized with respect to variance, remained in-tact. For these plots, we selected a maximum threshold of 1.6 SD above the mean BOLD signal because this was the largest value at which every brain region of every individual scan time series had a non-0 variance. Plot (d) illustrates this correspondence between the original and binarized example time series FC values of individual region pairs at a binarization threshold of 1, for which the correlation is 0.93. Plot (e), compares FC correlations between original and binarized time series data at binarization thresholds from 0 to 2.5. The median (black line) and range (shaded area) of individuals show that binarization reliably preserved individual FC up to a threshold of 1, whereas the FC correlation for the group aggregate data (red line) remained high up to a threshold of 2. At threshold 1, the worst case single-subject FC correlation was 0.88.

**Figure 1.**
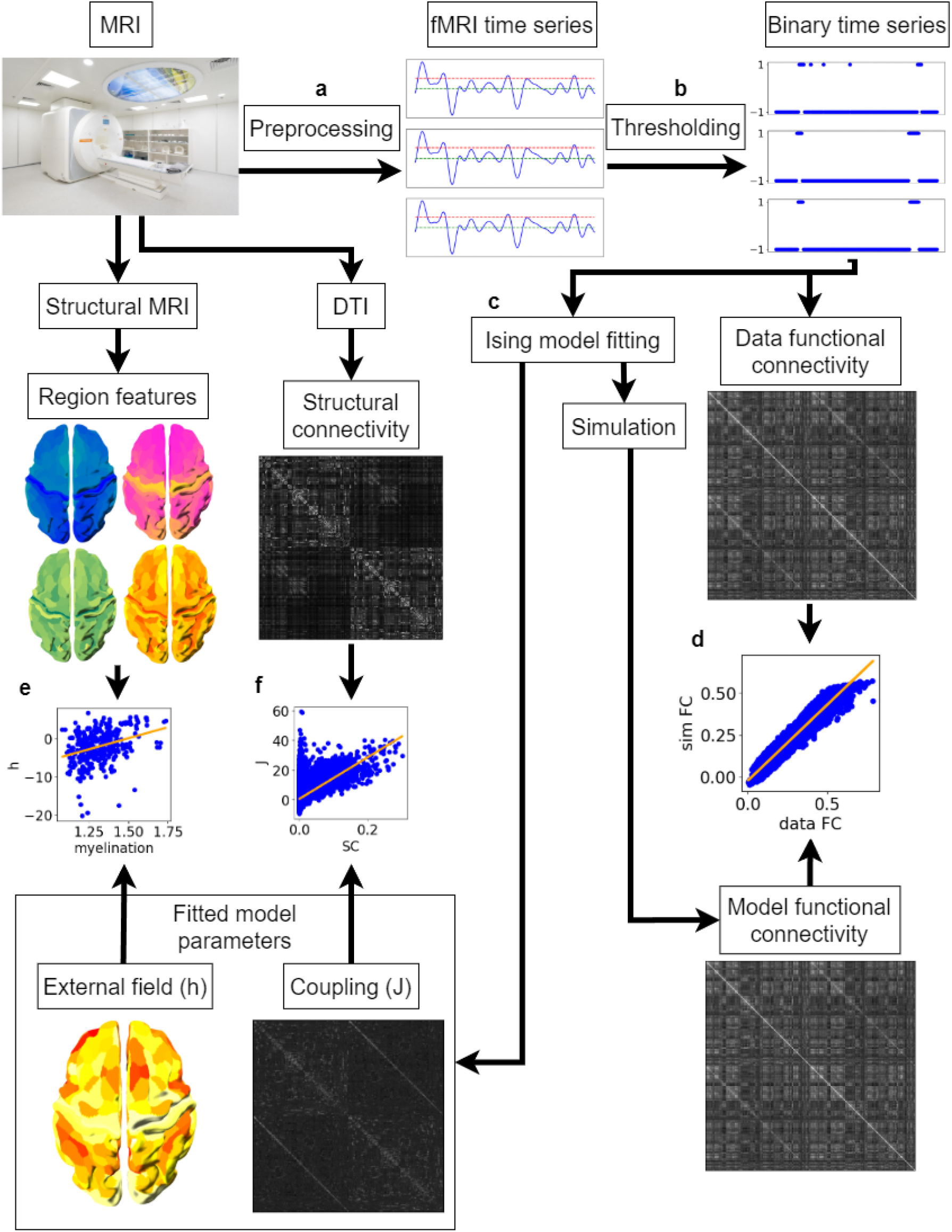
Ising model fitting and analysis workflow. (**a**) Preprocess fMRI data to get a continuous-value time series for each region. (**b**) Binarize the continuous-valued data (blue) using a threshold expressed in terms of standard deviations (red) above the mean (green) of the single region time series. (**c**) Optimize the model parameters, the external field of a region (*h*_*i*_) and coupling between regions (*J*_*ij*_), using Boltzmann learning, sampling states with the Metropolis algorithm. (**d**) Run a longer test simulation of the model to find the functional connectivity (FC). Then find the correlation between model and binarized data FC. (**e**) Find correlations between the fitted external field parameter value *h*_*i*_ and region structural features (thickness, myelination, curvature, and sulcus depth) from T1- and T2-weighted structural MRI. (**f**) Find the correlation between the coupling parameter *J*_*ij*_ and structural connectivity from DTI data.

**Figure 2.**
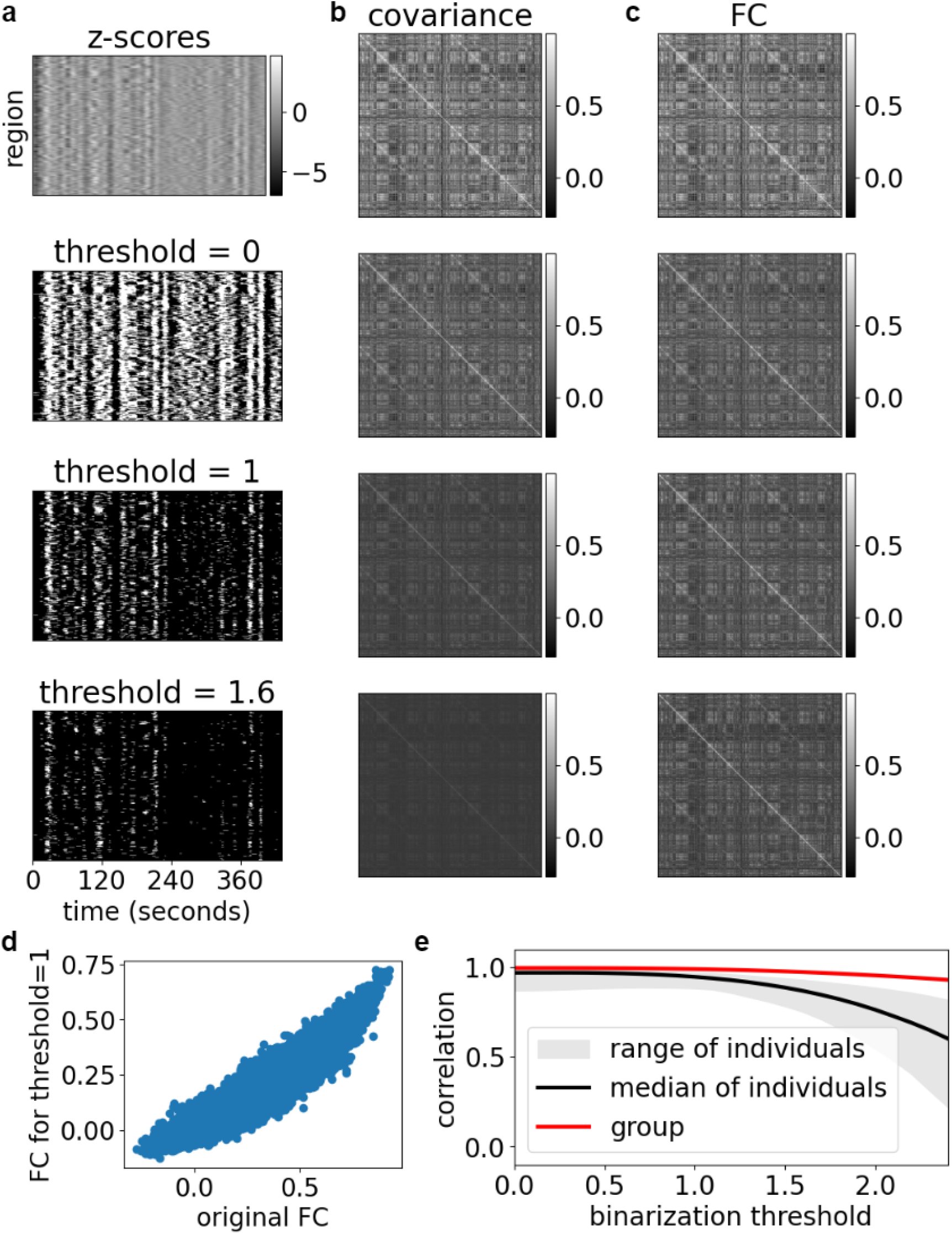
Effect of binarization on FC of fMRI time series data. (**a**) First half (432 seconds) of an example fMRI time series from a randomly selected HCP subject. The top image shows the preprocessed, parcellated, and z-scored but un-binarized values. Subsequent images show the same time series binarized at thresholds 0, 1, and 1.6 SD above the mean. (**b**) Covariances between pairs of regions in the original and binarized region time series. (**c**) FC (Pearson correlation) between pairs of regions in the original and binarized time series using all time points of all scans from the individual subject. (**d**) Scatter plot showing relationship between original and binarized FC of all region pairs for this subject. (**e**) Correlation between original and binarized time series FC as a function of binarization threshold. The black line shows the median, and the shaded area the range of individual subject FC correlations. The red line shows the FC correlation of group aggregate data (concatenated z-scored time series of 670 individuals).

**Figure 3:**
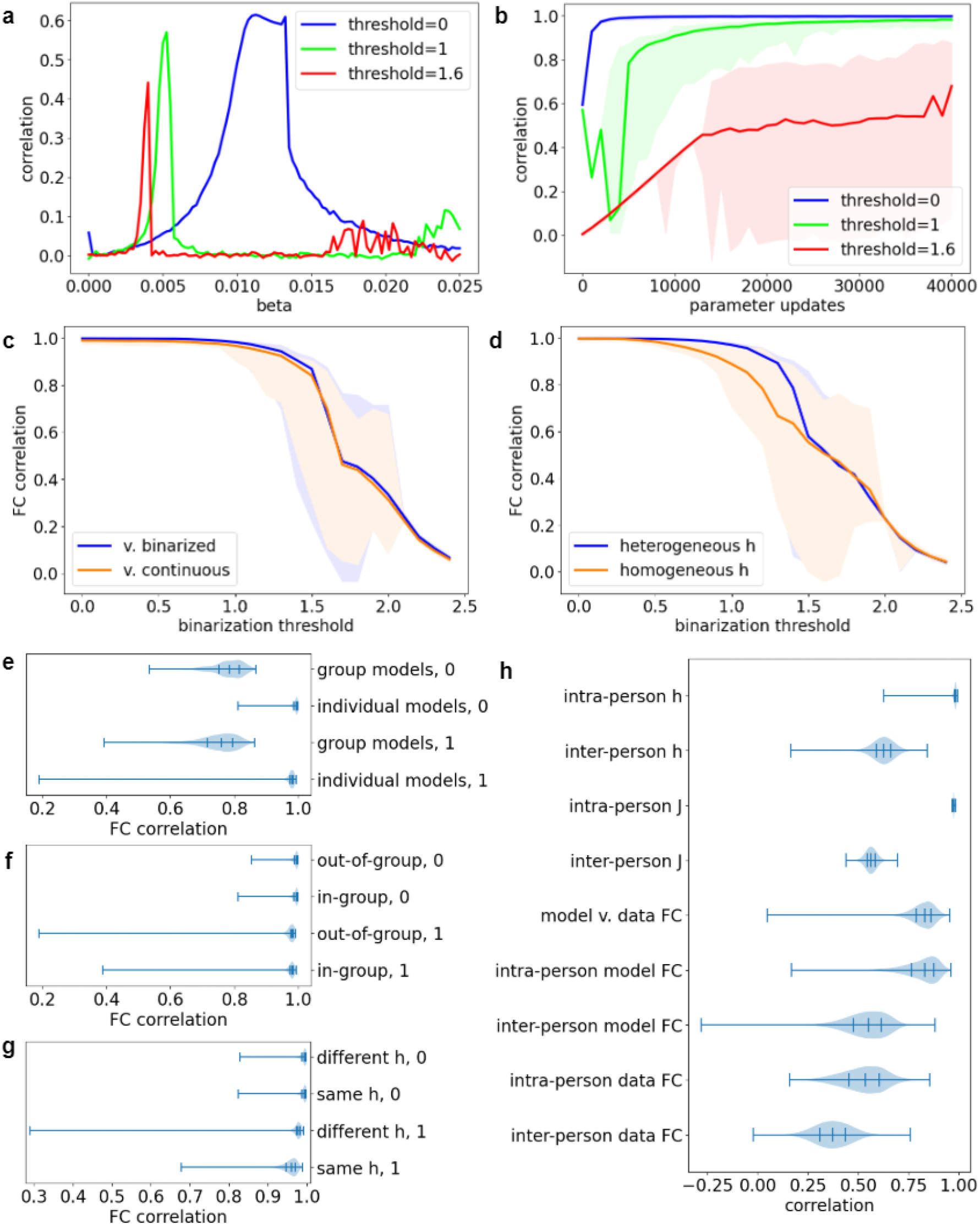
Goodness-of-fit measured by correlation between model and binarized data FC. (**a**) Effect of inverse temperature (beta) on correlation between the FC of binarized group-level data and the FC of the data-derived initial guess Ising model, prior to any fitting by Boltzmann learning. Each curve shows results for a different binarization threshold, 0, 1, or 1.6. (**b**) Progression of group model fitting by Boltzmann learning measured by correlation between binarized data FC and model FC. Lines represent medians, and shaded areas represent ranges over 101 independent fittings to the same data. (**c**) Correlations between FC of final fitted group models with either binarized group data (blue) or with continuous-valued group data (orange) at a range of binarization thresholds. The lines represent medians, and shaded areas represent ranges over 101 independently fitted replica models. (**d**) Correlation between the FC of the group model and binarized group data for the complete fitted model, including heterogeneous *h*_*i*_, (blue) and for a modified model using the same fitted value of *h*_*i*_ at all regions (orange). (**e**) Comparison of the distributions of group model FC versus individual data FC correlations and distributions of individual model FC versus individual data FC correlations. The number at the end of the label indicates the binarization threshold, 0 or 1. Ticks show the quartiles. (**f**) Comparison of distributions of FC correlations for individual models of subjects included in the group data fitting target (in-group) or excluded from (out-of-group) the group model fitting target. The number at the end of the label indicates the binarization threshold, 0 or 1. (**g**) Comparison of distributions of FC correlations for individual model versus individual data for full individual models, including fitted *h*_*i*_, and for individual models fitted with a modified method that allows individualized *J*_*ij*_ but forces all individuals to have the same *h*_*i*_ at each region. The number at the end of the label indicates the binarization threshold, 0 or 1. (**h**) Comparison of model and FC correlations for models fitted to single fMRI scans at threshold 1: region-wise correlations between elements of *h*_*i*_ for all pairs of different scans of the same person (intra-person h) or of two different people (inter-person h); region-pair-wise correlations between elements of *J*_*ij*_ for all pairs of different scans of the same person (intra-person J) or of two different people (inter-person J); region-pair-wise correlations between FC derived from simulating the fitted model and FC derived directly from the binarized data for single scans (model v. data FC); FC correlations for all pairs of Ising models fitted to different scans from the same person (intra-person model FC) or of two different people (inter-person model FC); correlations between pairs of FC matrices calculated directly from different scans of the same person (intra-person data FC) or from scans of two different people (inter-person data FC).

### 3.2. Goodness-of-fit of models fitted to group and individual data

Having determined the range of thresholds over which binarization preserves the FC of the data, we next tested our Ising model fitting workflow with the same data binarized at different thresholds. We also investigated the importance of the *h*_*i*_ parameters to model behavior. We found that, as the threshold increased, the initial guess model became less effective as an approximation of the fitted model, leading to slower and less reliable convergence, but, within the range where fitting succeeded, larger thresholds caused the heterogeneity of *h*_*i*_ play a more significant role in the simulation.

Figure 3a shows that, prior to fitting, each initial guess group model had an optimal simulation temperature that maximizes FC. However, as the threshold increased, the peak in FC correlation shrank and disappeared near 2.0. After 40,000 parameter updates, fitting converged to 0.98 or more for all models fitted to group data binarized at thresholds less than or equal to 1 but not at higher thresholds, such as 1.6 (Figure 3b). Past threshold 1, the final FC correlation dropped sharply (Figure 3c). Since binarization still preserved the group level FC in this range (see Figure 2e), FC correlation between the model and binarized data (blue line) matched FC correlation between the model and un-binarized data (orange line), even as both declined. Even though the differences in the threshold range [0, 1] were statistically significant (largest Wilcoxon *p* = 1. 7·10^−18^ < 1. 6·10^−3^), the largest difference between corresponding median-over-replicas FC correlations, which occurred at threshold 1, was 0.015. We also tested for convergence of the underlying model parameters and found that replica models remained nearly identical at thresholds up to 0.8 and remained mostly consistent except for some outlier replicas at higher thresholds (Supplementary Figure S7).

At thresholds 0 through 0.2, forcing *h*_*i*_ to be homogeneous across all regions during fitting had no effect (Wilcoxon *p* = 0. 48, 0. 039, *and* 0. 072, respectively), but, at thresholds 0.3 through 1.3, forcing homogeneity significantly worsened the fit between model and data (all Wilcoxon *p* < 4. 6·10^−8^; Figure 3d). At even higher thresholds, the effect of forcing homogeneity began to shrink again (Wilcoxon *p*∈[7. 2·10^−4^, 1. 0]). Specifically, at threshold 1, the median difference was 0.079, and *p* = 1. 7·10^−18^. We also found that, whereas the structure of the *J*_*ij*_ matrix remained similar and retained similar levels of heterogeneity at all thresholds, *h*_*i*_ vectors had similar patterns of heterogeneity across a wide range of positive thresholds but not at 0 (Supplementary Figure S8).

We next fitted models to individual data binarized at thresholds 0 and 1 and tested whether the individual variability of *h*_*i*_ played any role in reproducing individual differences in FC. While fitting succeeded at producing models with FC correlations at both thresholds in nearly all cases, threshold 1 performed worse in the worst cases. Whereas fitting at threshold 0 achieved FC correlations of 0.81 or more, fitting at threshold 1 yielded a small fraction of failed fitting attempts with FC correlations as low as 0.19. Of 4185 individual models fitted at threshold 1, 35 had FC correlations less than 0.90 (failure rate 0.0084). For threshold 0, the corresponding failure rate is 29 out of 4185 (0.0069). These failed fitting attempts did not appear to result from issues with any one subject’s data, as every subject had at least 2 replicas with FC correlations above 0.90. They did, however, appear to be genuine failures of model fitting instead of failures of the test simulation to converge to a stable FC, as the same replica models had low FC correlations across multiple tests. We also found that the FC correlations between individual models and individual data were slightly but significantly higher than the FC correlations between group models and individual data (Wilcoxon *p* = 6. 4·10^−139^ < 0. 025 for both thresholds; Figure 3e). FC correlations of the 670 subjects pooled for the group model, were slightly but significantly greater than those of the remaining 167 subjects (medians < 0. 001 apart for either threshold; 1-tailed Mann-Whitney U-test *p* = 2. 1·10^−89^ for threshold 0, *p* = 2. 3·10^−89^ for threshold 1, both < 0. 025; Figure 3f).

Forcing all individual models to have equal *h*_*i*_for a given region *i* did not result in significantly lower FC correlations at threshold 0 (1-tailed Wilcoxon *p* = 0. 22; Figure 3g) but did significantly lower FC correlations at threshold 1 (*p* = 1. 9·10^−130^ < 0. 025). We also examined the underlying model parameters and found that, *J*_*ij*_ of a given pair (*i, j*) had nearly the same mean and SD over individuals at either threshold. By contrast, mean *h*_*i*_ values did not correlate between thresholds 0 and 1, and the SD over individuals only weakly correlated while typically being twice as large for threshold 1 (Supplementary Figure S9), showing that the higher threshold had induced a greater degree and different pattern of heterogeneity among nodes.

To test how consistent models were across scans of the same individual, we fitted another set of Ising models to single resting-state fMRI scans binarized at threshold 1. We found that both *h*_*i*_ and *J*_*ij*_ were highly consistent between pairs of models fitted to different scans of the same individual. While the ranges of intra-person and inter person region-wise *h*_*i*_ correlations overlapped, only 9 of the 5022 intra-person pairs had correlations below the maximum inter-person pair *h*_*i*_ correlation (0.84). The distributions of region-pairwise *J*_*ij*_ correlations were even more distinct, with the range of intra-person correlations (0.97 to 0.99) completely disjoint from the range of inter-person correlations (0.44 to 0.70). For both comparisons, 1-sided Mann-Whitney U-test *p* = 0. See the first four violin plots in Figure 3h.

We then ran Metropolis simulations on the models to obtain a simulated FC for each. As with the 4-scan individual models at threshold 1, most models achieved high FC correlations with the fitting target (median 0.83, range 0.05 to 0.96, “model v. data FC” in Figure 3h). FC pairs generated from models of different scans of the same individual correlated more strongly than pairs from models of different individuals (1-sided Mann-Whitney U-test *p* = 0). The same held true for FC computed directly from the binarized data (*p* = 0). However, the medians of the model FC distributions were further apart (difference 0.28) than the medians of the data FC distributions (difference 0.16). Intra-person FC correlations were significantly higher for model-generated FC than for data FC (2-sided Wilcoxon signed-rank test *p* = 0) with only 32 model FC correlations less than the corresponding data FC correlations. See the last four violin plots in Figure 3h.

### 3.3. Predictability of model parameters from structural features

We compared parameters and structural features at two levels: brain-wide correspondences between group model parameters and group mean brain structural features, and local correspondences between individual differences in model parameters and individual differences in brain structure. We first computed the correlation between group model *J*_*ij*_ and the linear regression model *Ĵ* (*SC*_*ij*_) (Figure 4a, blue line). For comparison, we computed the correlation between *FC*_*ij*_ of the binarized group time series data and the linear regression model 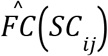 (Figure 4a, orange line). While *Ĵ* (*SC*_*ij*_) − *J*_*ij*_ correlation dropped and 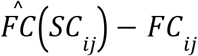 correlation rose as the threshold increased, both remained well above their permutation test critical values (all *p* < 10^−6^ < 8. 1·10 ^−4^), and *J*_*ij*_ correlation remained stronger at all thresholds.

**Figure 4:**
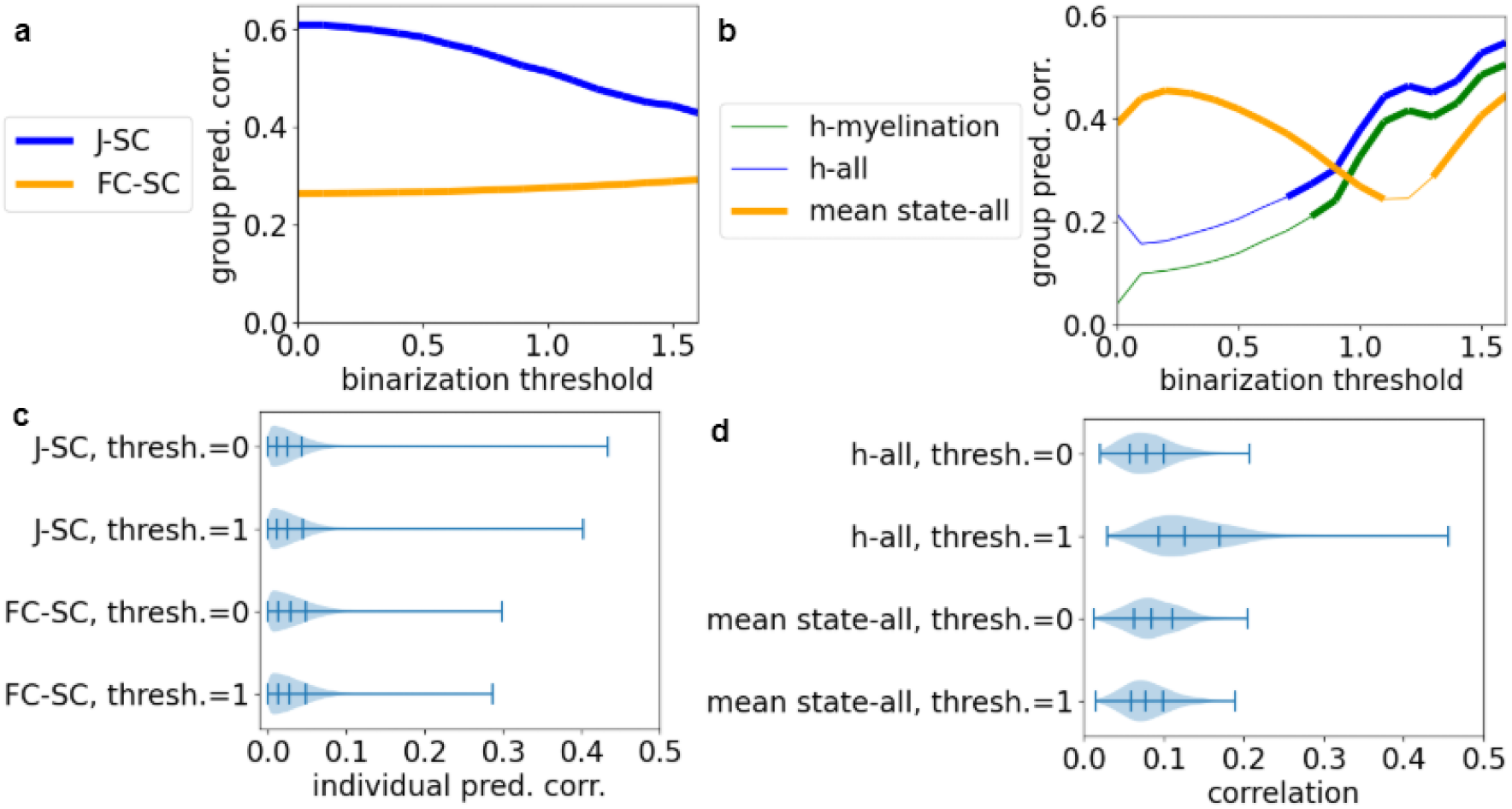
Correlations between fitted Ising model parameters and brain structural features. (**a**) Effect of binarization threshold on the correlation between group model *J*_*ij*_ and mean SC (blue) and the correlation between binarized group data FC and mean SC (orange). All correlations were significant (*p* < 8. 1·10^−4^). (**b**) Effect of threshold on correlations between group Ising model *h*_*i*_ and *h*_*i*_ as predicted from group mean structural MRI features (thickness, myelination, curvature, and sulcus depth) with a multiple least squares regression model (blue), *h*_*i*_ and *h*_*i*_ as predicted from mean myelination alone (green), and binarized mean state of the group-level data versus mean state predicted from the four structural features (orange). Thick line segments indicate significant correlations after Bonferroni correction (*p* < 1. 6·10^−4^). (**c**) Distribution of correlations between individual variations in individual Ising model *J*_*ij*_ and *J*_*ij*_ as predicted from SC with a least squares linear regression model (one linear model and correlation for each pair of brain regions) or between FC of the binarized data and FC predicted from SC. (**d**) Distribution of correlations between local variations in individual Ising model *h*_*i*_ and *h*_*i*_ as predicted from the four structural features with a least squares linear regression model (one linear model and correlation for each brain region)

As a further test of the robustness of these correlations, we cross-validated them using 10,000 random splits of region pairs into equal-sized training and testing sets. At all thresholds, training and testing correlations were largely identical, both forming a narrow band centered on the correlation for the full set of region pairs (Supplementary Figures S10 a and b). We also used the linear models that we fitted to group data to predict individual 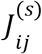 and 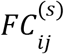 from 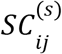 and found that, while the correlations (1 per subject) were lower, they were still consistently higher for *J*_*ij*_ than for *FC*_*ij*_ and were similar regardless of whether the subjects were included in the group model (Supplementary Figure S11a) or not (Supplementary Figure S11b).

We next compared group model *h*_*i*_ at multiple thresholds to group mean structural MRI features. To integrate information from all four features, for each threshold, we fitted a least squares multiple linear regression model *ĥ* (*t*_*i*_, *m*_*i*_, *c*_*i*_, *d*_*i*_) to *h*_*i*_, thickness (*t*_*i*_), myelination (*m*_*i*_), curvature (*c*_*i*_), and sulcus depth (*d*_*i*_). As expected, we found that the correlation between *h*_*i*_ and *ĥ* (*t*_*i*_, *m*_*i*_, *c*_*i*_, *d*_*i*_) trended upward with increasing threshold (Figure 4b, blue line), so that it first achieved statistical significance (indicated by a thicker line) at threshold 0.8 (correlation 0.27; *p* = 1. 4·10^−5^ < 1. 6·10^−4^). We observed the same upward trend in prediction correlation with threshold for the multiple least squares regression using the seven DTI and NODDI features (Supplementary Figure S14a).

To test the relative importance of different features, we compared these results to direct correlations with individual features and with single-feature linear regression models and found that *ĥ* (*t*_*i*_, *m*_*i*_, *c*_*i*_, *d*_*i*_)− *h*_*i*_ correlation closely tracked and was only marginally better than *ĥ* (*m*_*i*_)− *h*_*i*_ (Figure 4b, green line). The direct correlation between *h*_*i*_ and myelination was positive at all thresholds, while the other direct *h*_*i*_-feature correlations were negative when significant. All four became stronger with increasing threshold (Supplementary Figure 12a).

For comparison, we also computed the correlation between the mean state of the binarized group data, ⟨σ_*i*_ ⟩, and the linear model predicting it from structural features, 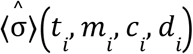 (Figure 4b, orange line). We found that 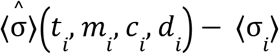 correlation changed non-monotonically with threshold, starting out higher than *ĥ* (*t*_*i*_, *m*_*i*_, *c*_*i*_, *d*_*i*_)− *h*_*i*_ correlation, dipping to a minimum near threshold 1.1, then rising again but remaining below *ĥ* (*t*_*i*_, *m*_*i*_, *c*_*i*_, *d*_*i*_)− *h*_*i*_ correlation. As with *ĥ* (*t*_*i*_, *m*_*i*_, *c*_*i*_, *d*_*i*_)− *h*_*i*_ correlation, the four-feature 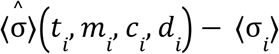 correlation closely tracked the single-feature 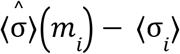 correlation, but, in this case, the direct *m*_*i*_ − ⟨σ_*i*_ ⟩ correlation flipped from negative to positive between thresholds 1.1 and 1.2 (Supplementary Figure S12b). Overall, the maximum correlations for *h*_*i*_ (0.55) and ⟨σ_*i*_ ⟩ (0.46) were close to each other.

We next tested whether individual differences in 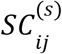 could predict individual differences in 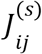 more accurately than they could predict individual differences in 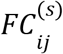. Figure 4c shows four distributions, each consisting of 64,620 correlations: 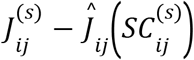 at threshold 0, 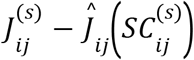 at threshold 1, 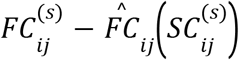 at threshold 0, and 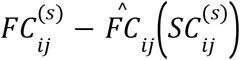 at threshold 1.

In all four cases, most of these local individual difference correlations were much weaker than the global correlation between group model and group mean data (Figure 4a), and fewer than 1% were statistically significant (1-tailed permutation test with Bonferroni-corrected α = 1. 9·10^−7^). 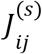 at threshold 1 had the most significant correlations (252), followed by 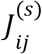 at threshold 0 (233), then 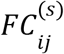 at threshold 0 (133), and finally 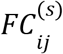 at threshold 1 (108). At either threshold, the overall size of the correlations was not greater for 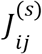 than for 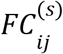 (both *p* = 1. 0). We confirmed the robustness of these findings by cross-validating with 100 random 670-167 (nearly 80-20) splits of the subjects into training and testing sets. While correlations that were weak on average tended to vary widely across different splits, the stronger correlations were more consistent, with mean training and testing correlations matching closely (Supplementary Figures S13 a-d).

Similarly, we tested whether local differences in structural features predicted local differences in *h*_*i*_. For each region *i*, we fitted a linear regression model 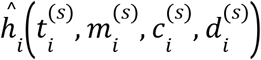 to individual Ising model 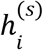 and structural feature values and computed the correlation between it and the Ising model 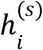 over all subjects *s*. We compared results for thresholds 0 and 1 and compared these to correlations between ⟨σ _*i*_⟩^(*s*)^ from the binarized data and linear regression models 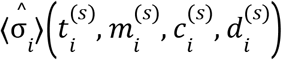. Figure 4d shows these four distributions.

Again, in all four cases, only a minority of regions had significant prediction correlations (Bonferroni-corrected α = 6. 9·10^−6^). 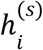 prediction correlations at threshold 1 were clearly higher than 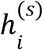 prediction correlations at threshold 0 (1-tailed Wilcoxon signed-rank test *p* = 1. 6·10^−39^) or ⟨σ _*i*_⟩^(*s*)^ correlations at either threshold 1 (*p* = 1. 3·10^−39^) or 0 (*p* = 1. 9·10^−31^) (Figure 4d). 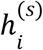 at threshold 1 also had the most regions with significant correlations, 59 compared to 2 for each other case. This was also true of direct correlations with individual structural features (Supplementary Figures S12 c-f). Unlike group-level *h*_*i*_, which mainly followed myelination, individual 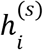 did not consistently have a single strongest predictor (Supplementary Figure S8h).

Cross-validation showed that, while weaker correlations were consistently lower over the testing set subjects than the training set subjects, this difference narrowed as the average correlations grew stronger (Supplementary Figures S13 e-h). For 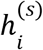 at threshold 1, individual difference prediction correlations for multiple least squares regressions using the seven NODDI and DTI features were generally weaker than those using the four T1- and T2-weighted structural MRI features (Supplementary Figure S14b).

### 3.4. Spatial trends in variability and predictability of Ising model parameters

We also searched for specific factors that could influence which connections had stronger predictability of individual 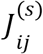 from 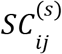 or which regions had stronger predictability of individual 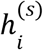 from structural features. For these analyses, we focused on individual Ising models at threshold 1 and compared prediction correlations to variability (SD) of structural features and model parameters. We found that 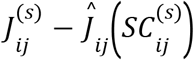 correlation moderately but significantly correlated with the SD of SC (0.37, 2-tailed permutation test *p* < 10^−6^, Figure 4e) and with SD of *J*_*ij*_ (0.25, *p* < 10^−6^).

Of the four structural MRI features, the 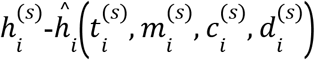 correlation correlated most strongly with SD of curvature (0.30, *p* < 10^−6^, Figure 4f). The correlations with the SD of thickness and SD of myelination were not significant (0.131, *p* = 0. 0085 and 0.064, *p* = 0. 109, respectively). While the correlation with SD of sulcus depth was significant (0.29, *p* < 10^−6^), the SD of curvature and SD of sulcus depth correlated strongly with each other (0.71), suggesting that they mostly provided redundant information. Cortical surface maps of 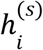 prediction correlation (Figure 4h) and of the SD of curvature (Figure 4i) show that, while the regions with the strongest structure-function couplings and those with the most variable curvature were not always the same, they tended to occur close to each other.

Comparing 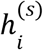 prediction correlations of regions and their descriptions in (Glasser et al., 2016) showed that the 59 regions with significant correlations occurred throughout the brain, but most occurred in the frontal and temporal lobes (Figure 5d). The broader area with the most regions with significant correlations was the anterior cingulate and medial prefrontal cortex (7 regions). Several other areas in the frontal lobe also had multiple significant correlations: the premotor cortex (5), inferior frontal cortex (4), orbital and polar frontal cortex (3), dorsolateral prefrontal cortex (2), and insular and frontal opercular cortex (2). Three of the temporal areas with the most significant correlations formed a single contiguous expanse: the ventral stream visual cortex (6), MT+ complex and neighboring visual areas (6), and lateral temporal cortex (6). Specifically, two of the significant regions (left and right PH) form a bridge between the ventral stream and the MT+ complex and border the lateral temporal cortex. The other parts of the temporal lobe with multiple significant correlations were the temporo-parieto-occipital junction (4), inferior medial temporal region (3), and auditory association cortex (2).

**Figure 5:**
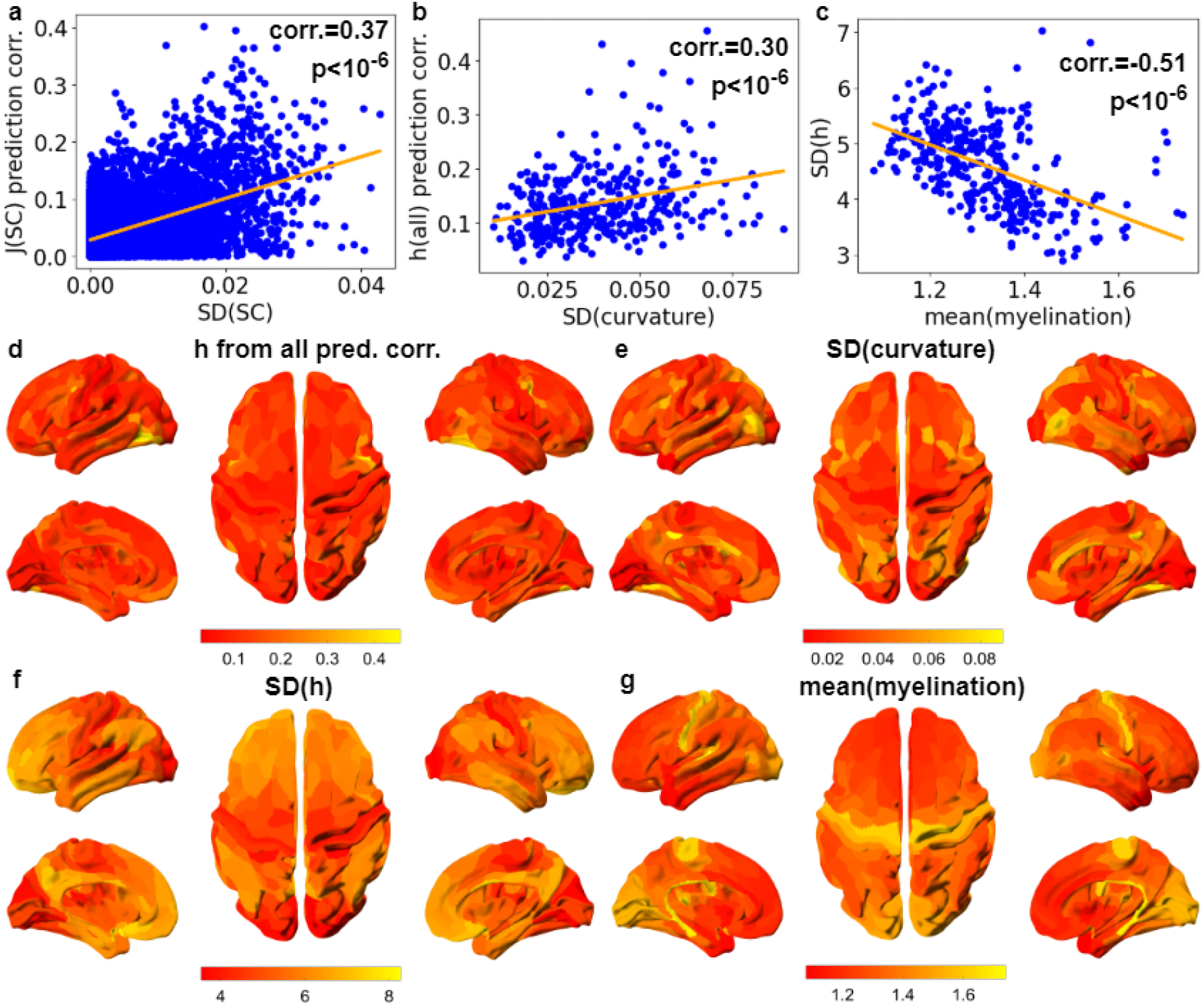
Spatial trends in variability and predictability of Ising model parameters. (**a**) Scatter plot of correlation between individual Ising model *J*_*ij*_ (threshold=1) and *J*_*ij*_ predicted from SC versus SD of SC (one point per pair of brain regions). The orange line shows the least squares regression trend. (**b**) Scatter plot of correlation between individual Ising model *h*_*i*_ (threshold=1) and *h*_*i*_ predicted from the four structural features versus SD of curvature (one point per brain region). The orange line shows the least squares regression trend. (**c**) Scatter plot of SD of individual Ising model *h*_*i*_ (threshold=1) versus mean myelination (one point per brain region). The orange line shows the least squares regression trend. (**d**) Cortical surface map of correlation between individual Ising model *h*_*i*_ (threshold=1) and *h*_*i*_ predicted from the four structural features. (**e**) Cortical surface map of SD of curvature. (**f**) Cortical surface map of SD of individual Ising model *h*_*i*_ (threshold=1). (**g**) Cortical surface map of mean myelination.

As a further test of the biological relevance of individual differences in 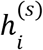. we tested whether more myelinated regions had lower variability of 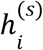. We first tested the correlation between 4-feature 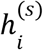 correlation and mean myelination. While the trend was positive, it was not significant (0.012, *p* = 0. 41 > 0. 0021). However, the local SD of 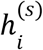 negatively correlated with mean myelination (−0.51, *p* < 10^−6^, Figure 4g). Specifically, the cortical maps of both mean myelination (Figure 4j) and SD of *h*_*i*_ (Figure 4k) show opposite trends along the axis from the highly myelinated sensory, motor, and occipital cortices to the less myelinated frontal and temporal areas.

## 4. Discussion

Overall, our work presents a more comprehensive approach to using Ising models as simplified, data-driven neural mass models to study the relationship between brain structure and functional characteristics. We started by introducing methodological improvements to enable Boltzmann learning to scale to larger, more fine-grained representations of the brain and achieve highly accurate fits to single-subject fMRI data. We then studied how data binarization threshold influenced the role of the heterogeneity of *h*_*i*_ in allowing the model to reproduce the FC of the data. Finally, we studied how binarization threshold affected relationships between *J*_*ij*_ and SC and between *h*_*i*_ and brain region structural features.

### 4.1. An improved approach to fitting to fMRI data

Fitting the group-level model by starting with a data-derived initial guess model, optimizing the simulation temperature prior to fitting the parameters, and using the group model as the initial guess for individual-level models lead to better convergence (Supplementary Figures S3 and S4). Whereas prior studies that fitted Ising models to fMRI data fitted them only to group-level data using pseudolikelihood maximization (Ezaki & Watanabe, 2017; Fortel et al., 2022; Ponce-Alvarez et al. 2022; Ruffini et al., 2023) or only fitted small models of 10 or fewer nodes to individual data using likelihood maximization (Watanabe et al., 2013; Theis et al., 2024; Theis et al., 2025) or Bayesian approximations (Jeong et al., 2021; Kang et al., 2021; Khanra et al., 2024), our approach can fit fully connected Ising models to a 360-region parcellation of individual-level fMRI data, allowing more precise identification of key regions and connections of importance to individual differences in patterns of whole-brain integration and segregation. As shown in Figure 3e, individualized models more accurately reproduced the FC of individual data than did group models. Furthermore, while correlations between individual differences in Ising model parameters and individual structural features were generally weaker than group-level spatial structure-parameter correlations, these correlations can serve as more sensitive measures of structure-function coupling than statistics calculated directly from the fMRI data (Figures 4c, 4d).

Furthermore, our study is the first to address how choice of binarization threshold affects the role of node heterogeneity in the Ising model, which is important to comparisons between the Ising model and the real brain, in which brain regions have distinct intrinsic dynamical properties. Of the prior studies that used fMRI data as a fitting target, only (Theis et al., 2025), (Fortel et al., 2022), and (Watanabe et al., 2013) compared Ising models fitted to data binarized at multiple thresholds. All three reported that binarization at the mean produced the closest fit between model and data. While this also held true for our method, it achieved good correlations among model simulation FC, binarized data FC, and continuous-valued data FC over a range of thresholds from 0 to 1 SD above the mean BOLD signal.

Whereas the above-cited works did not systematically examine the effects of binarization threshold on the underlying parameters of the fitted model, we showed that it influenced the extent to which the model was merely a network of interchangeable nodes or a network of nodes with distinct functional characteristics. At a threshold of 0, removing region-to-region heterogeneity of *h*_*i*_ did not affect the FC correlation of the group model, and removing subject-to-subject heterogeneity of *h*_*i*_ did not affect the FC correlation of the individual models. This implies that the nodes were effectively homogeneous and that *J*_*ij*_ contained all information needed to reproduce the FC of the data. By contrast, using a threshold of 1 maintained good FC correlation and induced both the group and individual models to use heterogeneous *h*_*i*_ values that played significant roles in reproducing the FC of the original data. This better reflects the real human brain, in which distinct regions have different morphology and in which individual differences in local region structural features correlate substantially with excitability (Glasser et al., 2014; Fotiadis et al., 2023; Fotiadis et al., 2024), which in turn influences larger-scale brain dynamics (Abeywardena, 2024).

We focused on models with a threshold of 1 as this was the highest threshold at which FC correlation remained strong, allowing us to demonstrate the maximal effect of regional heterogeneity. However, we have also shown in supplementary section S8 that group model parameters remain strongly correlated with each other across a wide range of thresholds with 0 and other very low thresholds representing a special case where *J*_*ij*_ remains correlated with versions of it for higher thresholds but *h*_*i*_ does not correlate with *h*_*i*_ at other thresholds and has low variance. This suggests that future studies that use this approach to Ising model fitting can adjust the threshold to reflect the relative importance of regional heterogeneity and connectivity to the research question. For instance, if the project requires more accurate reproduction of the data FC, a lower non-0 threshold can achieve this while still preserving significant regional heterogeneity. At the other end of the range, examinations of avalanche criticality in fMRI data, such as (Wang, 2019), may use thresholds close to 1.5 or even 2. An Ising model fitted to this same binarized data would only poorly reproduce the FC but could still have parameters that correlate significantly with structural features.

Furthermore, when we fitted models to single fMRI scans, we found that, not only were model-generated FC correlations significantly more similar for different scans of the same individual than for scans of different individuals, but the fitted model parameters were highly consistent between different scans of the same person (Figure 3h). This shows that our fitting method allows Ising models to capture persistent, distinctive features of an individual person’s brain activity. Furthermore, intra-person FC correlations were higher for model-generated FC than for FC computed directly from the binarized data (Figure 3h, last 4 violin plots). This suggests that, while the model accurately captures an individual’s static FC, it does not necessarily capture the full variability of their FC over shorter windows.

### 4.2. An intermediary between structural and functional connectivity

To test the biological relevance of the fitted model parameters, we computed correlations between *J*_*ij*_ and SC and between *h*_*i*_ and four local region features: thickness, myelination, curvature, and sulcus depth. We found that, for any FC-preserving binarization threshold, the least squares regression model prediction from group mean SC correlated better with group model *J*_*ij*_ than with group binarized data FC, showing that the Ising model can serve as an intermediary between the structural network and the functional network (Figure 4c). As suggested in (Ruffini et al., 2023), *J*_*ij*_ could serve as a signed approximation of the structural connectome in future modeling studies with more sophisticated models such as neural mass models, where recovering the signed structural connectivity from functional data is more difficult.

While earlier studies have found similarly robust relationships between population-level SC and *J*_*ij*_ (Watanabe et al., 2013; Fortel et al., 2022; Ruffini et al., 2023), our approach allowed us to study the relationship between individual differences in local SC and *J*_*ij*_ in a way that other studies have not. While individual-level correlations were much more variable and generally much weaker, the number and strength of significant SC-*J*_*ij*_ correlations were significantly greater than for FC-SC correlations (Figure 4c and Supplementary Figure S12g). Furthermore, the largest SC-*J*_*ij*_ correlations occurred on connections with greater variability of SC (Figure 5a), suggesting that the individual Ising models do succeed at capturing the most significant differences in underlying connectivity. These results indicate that, in future studies, the *J*_*ij*_ can serve as an intermediate representation through which to link individual differences in fMRI-derived FC and DTI-derived SC more effectively than direct correlations between the two.

### 4.3. A mapping from structural features to external field

Whereas researchers have previously noted the correspondence between SC and *J*_*ij*_ (Watanabe et al., 2013; Ponce-Alvarez, 2022; Ruffini et al., 2023), none have studied the relationship between *h*_*i*_ and brain structure. Whereas mean activation of a region depends on both its excitation-inhibition (E-I) balance and its connectivity with other regions, the Ising model fitting process disentangles these aspects of local behavior, capturing connectivity in *J*_*ij*_ and the probability that a region will be differentially activated, given the same inputs, in *h*_*i*_. The correlation between myelination and *h*_*i*_, considered in the context of prior studies relating myelination to E-I balance (Glasser et al., 2014; Fotiadis et al., 2023; Fotiadis et al., 2024), suggests that it reflects the overall excitability of the local region, though stronger conclusions about the interpretation of *h*_*i*_ will require future studies comparing it to more direct measurements of the intrinsic excitability, possibly through transcranial magnetic stimulation, as used in (Roos et al., 2021).

That group-mean T1w/T2w ratio correlated strongly with group model *h*_*i*_ suggests that this method of measuring myelination is adequate for finding the general orientation of the principal axis along which *h*_*i*_ varies, but the even stronger group-level correlations that we obtain with DTI and NODDI features (Supplementary Figure S14b) suggest that combining information from multiple myelin-associated features can help to more accurately predict the functional gradients along which *h*_*i*_ varies. However, whereas T1w/T2w ratio correlates positively with *h*_*i*_, NDI correlates negatively, suggesting that the two measures follow different spatial gradients.

Combining multiple such features by using a multiple linear regression model produced better predictions of *h*_*i*_, showing that multiple aspects of brain structure influence activity. As with the SC-*J*_*ij*_ correlations, we considered both region-to-region heterogeneity based on pooled population-level data and local individual differences. While group-level *h*_*i*_ prediction correlations were robust, they were only slightly stronger than the best group-level binarized mean state prediction correlations (Figure 4b). By contrast, *h*_*i*_ clearly achieved stronger correlations with individual differences in structure (Figure 4d). This suggests that *h*_*i*_ is especially valuable for representing individual rather than group differences in excitability and can serve as a more sensitive measure of structure-function coupling.

To develop a better sense of how specific structural features influenced these differences, we searched for region-to-region trends in structural features that correlated with the individual-level variability or predictability of *h*_*i*_. At the whole-brain level, myelination was the dominant predictor of group-level *h*_*i*_ (Supplementary Figure S12a), which increased along the axis from less myelinated multimodal information integration regions to unimodal sensory and motor regions. This is consistent with the hypothesis that myelination offers increased efficiency of signal transmission, represented in the Ising model by easier induction of differential activation due to higher *h*_*i*_. However, along this same axis, we saw a decrease in individual variation in *h*_*i*_ (Figure 5c), suggesting this increased efficiency comes at the cost of reduced synaptic plasticity. While prior studies such as (Paquola, 2019) and (Fotiadis et al., 2023) have argued for this tradeoff based on mechanistic hypotheses and correlations between group-level functional and structural gradients, our results show that *h*_*i*_ can provide a measure of both spatial group-level functional gradients and individual-level differences derived from fMRI data by disentangling excitability (measured by *h*_*i*_) from connectivity (measured by *J*_*ij*_).

However, the variability of curvature, not myelination, had the strongest relationship with the predictability of *h*_*i*_ (Figure 5b). Furthermore, combining information from multiple features offered a substantial advantage when predicting individual differences in *h*_*i*_ (Compare Figure 4d to Figures S12 c-f). This suggests that, whereas myelination constrains the capacity for individual variation, the actual differences in brain dynamics arise through changes in gray matter cortical architecture, particularly through differences in the amount of cortical folding. This is consistent with an earlier study, which found that sulcus depth correlated with individual variation in the resting state functional connectivity between a region and surrounding areas (Mueller et al., 2013). That multiple least-squares regression using the combined DTI and NODDI features, which include six features, all associated with myelination and other aspects of microstructure, resulted in lower prediction correlations than those achieved with the T1- and T2-weighted structural MRI features (Supplementary Figure S14c) further supports this interpretation. That the highest *h*_*i*_ prediction correlations concentrated in the frontal lobe and in higher visual and multimodal areas also suggests that regions with higher synaptic plasticity have more room for changes in architecture that can influence *h*_*i*_.

### 4.4. Limitations and future work

The present work has four key limitations: the simplicity of the Ising model itself, the consideration of only local, linear relationships between Ising model parameters and structural features, the consideration of only resting state brain activity, and the consideration of only healthy subjects.

We can only fit the Ising model to the mean activation and pairwise covariances of the data, which do not capture its temporal dynamics (see Methods). However, our research group is currently working to develop a mathematical framework for relating Ising model parameters to continuous-valued dynamical models. A possible intermediate step is the generalization of our approach from symmetric Ising models that generate a static distribution of states to asymmetric, kinetic Ising models, which have binary node states but can capture dynamically varying covariance (Zeng et al., 2013).

The present work only relates the *h*_*i*_ value of a region to its structural MRI features and the *J*_*ij*_ value of a pair of regions to the SC between them. This approach assumes that the the Ising model parameters reflect only the influence of local features and that these influences are linear. That SC-*J*_*ij*_ correlation is stronger than SC-FC correlation suggests that *J*_*ij*_ has a stronger coupling to local SC than does FC with less influence from indirect connectivity, but the structural influences on *J*_*ij*_ and *h*_*i*_ may still not be fully local. Incorporating indirect structural connections improves prediction of FC (Røge, 2017), so the same may improve prediction of *J*_*ij*_. Additionally, the local relationships between structural features and dynamical properties may be nonlinear. In particular, (Fotiadis, 2023) makes the case that myelination and excitation-inhibition balance interact synergistically to influence structure-function coupling, which varies widely between highly myelinated sensory and motor regions and less myelinated associative areas. The authors of the present work are currently comparing different machine learning approaches that can detect non-local and non-linear relationships between Ising model parameters and structural features.

Another open question is the relationship between individual differences in Ising model parameters and individual differences in cognitive abilities. In the present work, we show that models can capture features of individual resting-state brain activity that persist across scans, but this does not imply that we can predict how they will change when transitioning to task conditions. Relating these regional and pairwise features to a system-level property of human behavior requires a very different methodology from what we have used to map local structural features to local Ising model parameters. Our research group is applying parameter sensitivity analysis of Ising models fitted to both resting and task condition fMRI data to identify directions in parameter space that both control meaningful differences in model FC and correlate with differences in participants’ working memory task performance, providing a new approach to defining task-related subnetworks (Chen et al., 2025).

Finally, we have only fitted our model to data from healthy young adults in the absence of any neuromodulation, meaning that our model cannot currently predict the effects of maturation, aging, neurological disorders, or medical interventions. Members of our research group are currently applying the approaches used here and in (Chen et al., 2025) to compare how Ising model parameters change during maturation in both autism spectrum disorder patients and neurotypical controls. The goal of this work will be to identify the local and network-level differences in brain structure that most strongly influence differences in both brain activity and behavioral phenotype, which can then serve as putative targets for novel therapies.

## Conclusion

The Ising model fitting workflow and parameter analyses presented here represent a substantial step forward in the use of Ising models as simplified, data-driven neural mass models. While fitting a high-dimensional model using Boltzmann learning remains computationally costly, our implementation decreases the time needed to fit an ensemble of models by fitting them in parallel on a GPU. By fitting models to whole-brain data instead of pre-selecting a set of regions to include, researchers can discover important network-level individual differences. Furthermore, selecting a threshold that induces node-level heterogeneity will produce models that reflect both the interconnectivity and distinctness of brain regions. This approach will help to bridge the gap between connectomics and translational neuroscience. While connectomics draws on increasingly complex network-level views of the brain (Mišić & Sporns, 2016), clinical research continues to focus on single regions when designing interventions such as transcranial direct current stimulation (Kang et al., 2024) and transcranial magnetic stimulation (Cash & Zalesky, 2024). Future work will focus on both the discovery of nonlinear and non-local associations between structural features and Ising model parameters and the identification of networks of regions and connections that are important to individual differences in cognitive abilities and mental health.

## Supporting information

Supplementary analyses of alternative methods and additional validations

## ACKNOWLEDGMENTS

This work was partially supported by STI 2030-Major Projects (No. 2022ZD0208500), the Hong Kong Research Grant Council Senior Research Fellow Scheme (SRFS2324-2S05) and General Competitive Fund (GRF12202124), and Hong Kong Baptist University (HKBU) Seed Funding for Collaborative Research Grants (RC-SFCRG/23-24/SCI/06). Data were provided by the Human Connectome Project, WU-Minn Consortium (Principal Investigators: David Van Essen and Kamil Ugurbil; 1U54MH091657) funded by the 16 NIH Institutes and Centers that support the NIH Blueprint for Neuroscience Research; and by the McDonnell Center for Systems Neuroscience at Washington University.

The authors thank Zhao Chang of the HKBU Department of Physics for his help in using the AMICO pipeline and interpreting its output.

## Author contributions

Conceptualization: CZ; Methodology: AC, SC, QT; Investigation: AC; Visualization: AC, SC, QT; Writing—original draft: AC; Writing—review \& editing: all authors.

## Data and materials availability

The MRI data are available at https://www.humanconnectome.org/study/hcp-young-adult. The code used for this study is available on request.

## Data ethics statement

All data used in this study come from the Human Connectome (HCP) Project 1200 Subjects data release. Our use of the data complies with all terms of use available at https://humanconnectome.org/study/hcp-young-adult/document/wu-minn-hcp-consortium-open-access-data-use-terms (last accessed 2025-10-21). Researchers contributing data to the dataset first obtained consent from participants as described in the HCP S1200 standard operating procedures: https://humanconnectome.org/storage/app/media/documentation/s1200/HCP_S1200_Release_Appendix_IV.pdf. We do not believe that it is possible to use the data and statistical aggregates presented in this article to identify any individual HCP participant.

Specifically, we only use functional and structural brain imaging data (fMRI, DTI, T1-weighted and T2-weighted structural MRI) and do not use the Restricted HCP Data described at https://humanconnectome.org/study/hcp-young-adult/document/restricted-data-usage (last accessed 2025-10-21).

