## Supplementary analyses of alternative methods and additional validations for "Personalized whole-brain Ising models with heterogeneous nodes capture differences among brain regions"

### S1. Fitting Ising models to AAL-parcellated fMRI data

To test whether our fitting method could generalize to other parcellations of the brain, we parcellated the 4 fMRI time series per subject from the 837 selected subjects using the 117-region Automatic Anatomical Labeling (AAL) Atlas Version 1 (Tzourio-Mazoyer et al., 2002) and pooled the same 670 of them to fit a set of group models using the same approach as with the 360-region Glasser Atlas (Glasser et al., 2016). We tested 31 thresholds evenly spaced between 0.0 and 3.0, inclusive of both endpoints. Figure S1a shows that, at each threshold, the data-derived initial guess model attains a higher maximum FC correlation and remains close to that correlation over a wider range of $\beta$ compared to the Glasser Atlas (Figure 3a in the main text). We then fitted 101 replica models to each target derived from binarized group data at a different threshold using Boltzmann learning for 30,000 parameter updates, following the same procedure as with the Glasser Atlas parcellated data. We again found that the FC correlation between model and binarized data drops sharply as the binarization threshold exceeds 1 (Figure S1b and S1c). We also tested the effect of replacing each $h_i$ parameter with the mean over all regions and found that this only affected FC correlation between the model and binarized data at thresholds above 0 (Figure S1c). For each of the 837 subjects, we then used all four scans binarized at a threshold of either 0 or 1 to create an individual data fitting target and fitted 5 replica models to it, using the best group model for each threshold as the initial guess. We then compared FC correlations for these fitted models to those of modified versions with $h_i$ replaced with the mean over all subjects of $h_i$ for the same region (Figure 3d). While a 1-tailed Wilcoxon signed rank test did show that removing individual differences in $h_i$ did significantly decrease FC correlation for threshold 0 ($p < 10^{-8}$), the differences were consistently small, the largest being 0.00152. By contrast, for threshold 1, removing individual differences in $h_i$ consistently and dramatically degraded FC correlation. The smallest difference was 0.0997, about 10% of the higher correlation value. Overall, these results show that, while fitting to lower-dimensional data parcellated with the AAL Atlas is slightly faster, our method still yields similar FC correlations for both Atlases, and the use of a non-0 binarization threshold is still key to inducing heterogeneity among the Ising model nodes to play a significant role in model behavior.

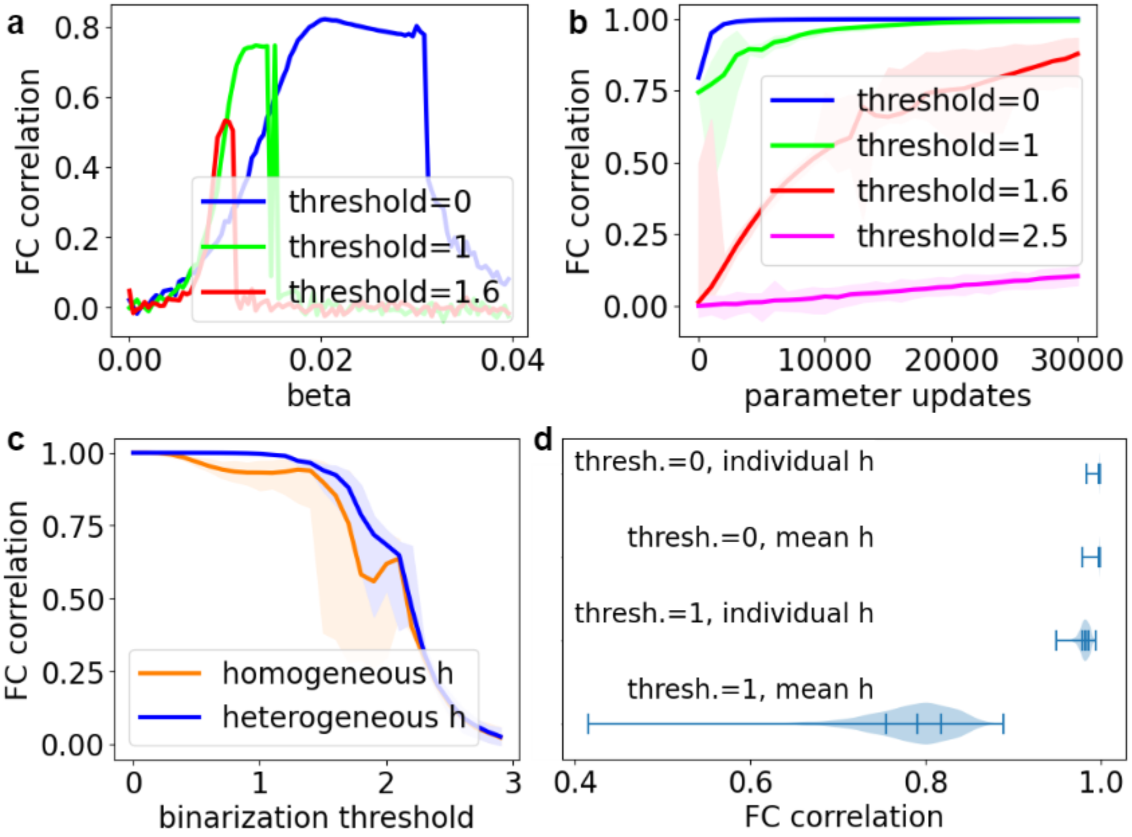


**Figure S1. Fitting Ising models to AAL-parcellated data.** (**a**) FC correlation of the AAL-parcellated-data-derived initial guess group model as a function of $\beta$. For each binarization threshold, we tested the initial guess model derived from the pooled group data parcellated with the AAL Atlas by simulating for 120,000 steps and then comparing the simulation FC to that of the binarized data. (**b**) Fitting progress of AAL group Ising models. For each binarization threshold, we fitted 101 replica models for 30,000 parameter updates. Lines represent medians, and shaded areas represent ranges. (**c**) Comparison of FC correlation of group models after fitting (blue) compared to modified versions with $h_i$ replaced with the mean value over all regions $i$. (**d**) Comparison of the distributions of FC correlations of individual models of 837 subjects with modified versions with the individual $h_i$ replaced with the mean $h_i$ for the same region over all subjects. We fitted models to convergence to individual data binarized at threshold 0 or threshold 1. Tick marks represent quartiles.

### S2. Fitting with pseudolikelihood maximization

Pseudolikelihood maximization is another commonly used method of fitting Ising models to data. While potentially less accurate, it is usually less computationally expensive than Boltzmann learning (Ezaki & Watanabe, 2017). As with Boltzmann learning, pseudolikelihood maximization consists of iterative updates to the model parameters determined by the differences between the empirical means (for $h_i$) and uncentered variances (for $J_{ij}$) of the data and counterparts computed from the model:

$$\delta h_i = \eta\left(\langle\sigma_i\rangle_{\text{data}} - \langle\sigma_i\rangle_{\text{pl}}\right), \delta J_{ij} = \eta\left(\langle\sigma_i\sigma_j\rangle_{\text{data}} - \langle\sigma_i\sigma_j\rangle_{\text{pl}}\right) \text{ (S1)}$$

However, instead of running the Metropolis simulation to sample states from the model and compute the model state means and uncentered covariances, this approach calls for estimation of the first- and second-order mean fields, $\langle\sigma_i\rangle_{\text{pl}}$ and $\langle\sigma_i\sigma_j\rangle_{\text{pl}}$, from the current model parameters and the time series data:

$$\langle\sigma_i\rangle_{\text{pl}} = \frac{1}{T}\sum_{t=1}^{T} \tanh\left( h_i + \sum_{j=1, j\neq i}^{N} J_{ij}\sigma_j\,(t) \right) \text{ (S2)}$$

$$\langle\sigma_i\sigma_j\rangle_{\text{pl}} = \frac{1}{T}\sum_{t=1}^{T} \sigma_i(t) \tanh\left( h_i + \sum_{j=1, j\neq i}^{N} J_{ij}\sigma_j\,(t) \right) \text{ (S3)}$$

Here, $\sigma_i(t)$ is the state of region $i$ at time point $t$ of $T$ in the data (*ibid.*). These formulae come from the pseudolikelihood, an approximation of the likelihood of the model parameters given the data:

$$\mathcal{L}(\boldsymbol{h}, \boldsymbol{J}) \approx \prod_{t=1}^{T}\prod_{i=1}^{N} P(\sigma_i(t)|\boldsymbol{h}, \boldsymbol{J}, \sigma_{/i}(t)) \text{ (S4)}$$

Here, $\sigma_{/i}(t)$ is the set of all spins of regions other than that of region $i$ fixed to their values at time point $t$ in the data, and $P$ is the Boltzmann distribution (Equation 1 in the main text) (*ibid.*).

We used the pseudolikelihood method from (*ibid.*) to fit Ising models to the 670-subject pooled group HCP fMRI data binarized at thresholds 0, 1, and 2.4. We used the same initial guess models and learning rate (0.01) as we used when fitting with Boltzmann learning but fitted for up to 300,000 parameter updates instead of only 40,000. We found that the models at 80,000 updates and 300,000 updates were effectively identical, indicating that fitting had converged.

Because pseudolikelihood maximization is deterministic, we only fitted one group model per threshold. We then simulated each fitted Ising model for 120,000 steps at a range of $\beta$ to find the one that achieved the best FC correlation. Figure S2 compares performance at the selected thresholds. For threshold = 0 (blue line), the optimal $\beta$ is 0.84, and it produces an FC correlation of 0.982, only slightly worse than what Boltzmann learning achieves, 0.9857 to 0.9992. For threshold = 1 (green line), the best $\beta$ is 0.65, and it achieves an FC correlation of 0.755, substantially worse than the final FC correlations for Boltzmann learning, 0.9452 to 0.9888. For threshold = 2.4 (red line), the best $\beta$ is 0.62, and it achieves an FC correlation of 0.745, whereas Boltzmann learning completely fails, with final FC correlations ranging from 0.0567 to 0.0808. Overall, pseudolikelihood maximization is less effective than Boltzmann learning in the range where binarized FC is close to that of the original data but may be useful for fitting an Ising model to data binarized at more extreme thresholds when a loose approximation is sufficient.

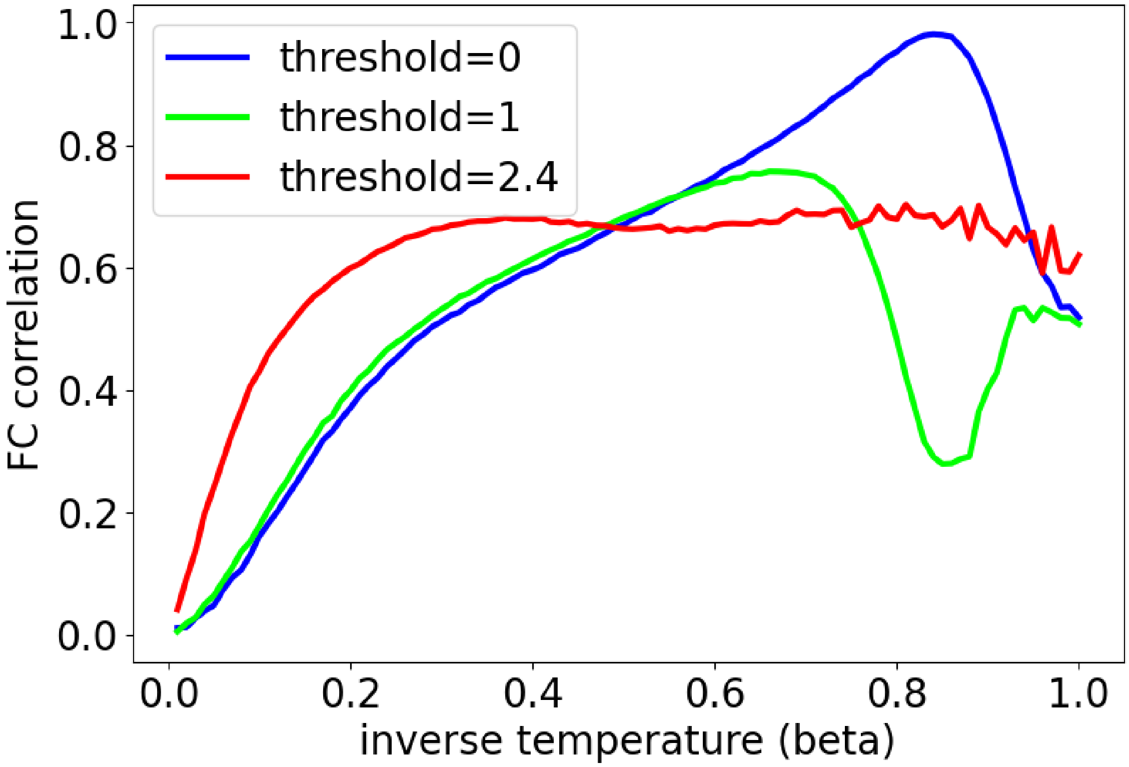


**Figure S2. FC correlations of group models fitted by pseudolikelihood maximization.** For each binarization threshold, we binarized each time series, concatenated the binarized time series into a group data set, initialized the model using the group mean and uncentered covariance, fitted the model by pseudolikelihood maximization for 300,000 parameter updates, performed 120,000 steps of Metropolis simulation at each of 101 values of $\beta$, and computed the correlation between the model and data FC at each (threshold, $\beta$) pair.

### S3. Randomized initial parameters

To demonstrate the value of our data-derived initial parameters, we compare to a model initialized with parameters sampled from normal distributions with the same means and variances as our data-derived initial parameters for the group model. That is, the external fields have the same mean and variance as the distribution of mean states, and the couplings have the same mean and variance as the distribution of uncentered covariances of pairs of states. We repeat this initialization for each of three binarization thresholds: 0, 1, and 1.6 SD above the mean. We then test the random models at 101 different inverse simulation temperatures ($\beta$). Unlike our data-derived initial Ising models, the simulation FC values from the randomly generated models do not show a clear correlation with the corresponding data-derived FC at any temperature or binarization threshold (Figure S3a). The range of $\beta$ tested covers the phase transitions of all models, as we can see by plotting the flip rates (Figure S3b). That the FC of the random model does not correlate with the data FC at any $\beta$ shows that the correlation between the simulated FC of the data-derived initial guess model and binarized data FC at optimal temperature is non-trivial. We then compare how quickly fitting converges for the data-derived and randomized initial guess models using the same simulation temperatures. The FC correlations quickly converge toward similar ranges, suggesting that the benefit of using a data-derived initial guess model is that it enables selection of a suitable temperature for the choice of binarization threshold, in this case 0 (Figure S3c) or 1 (Figure S3d).

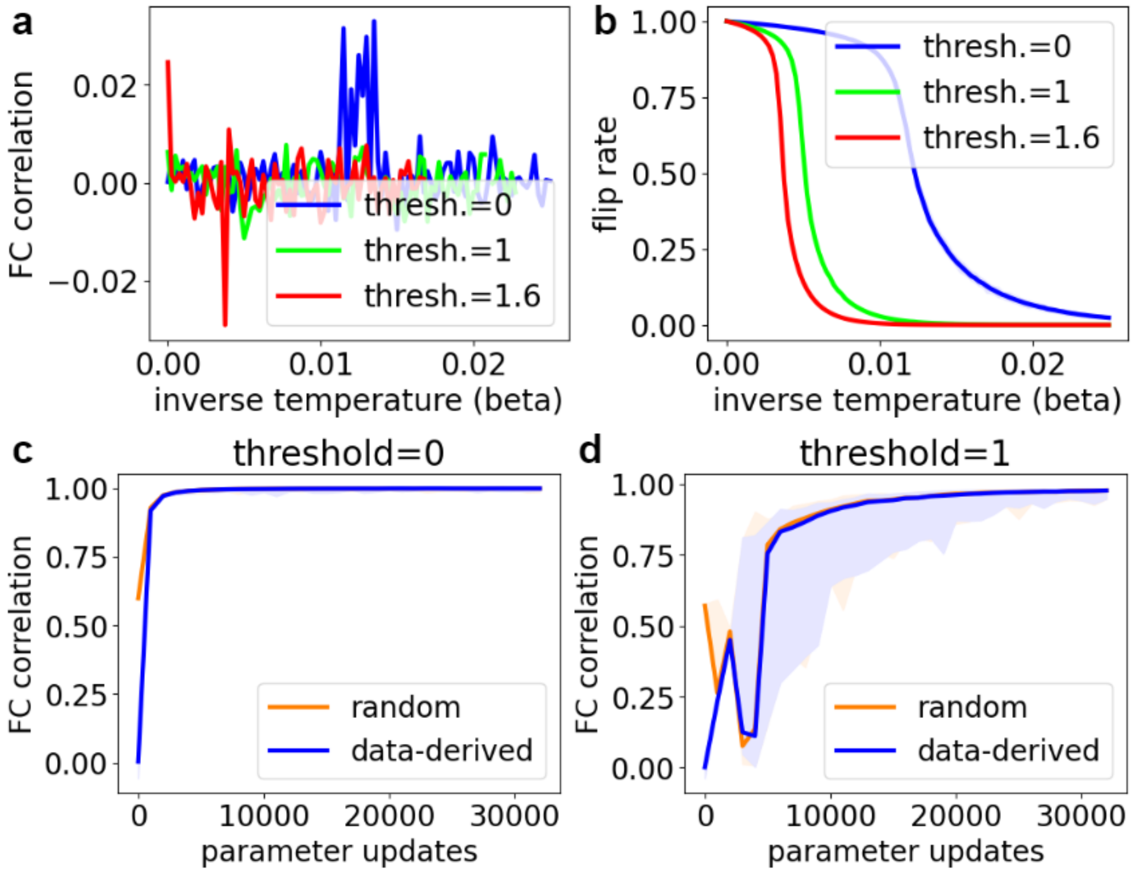


**Figure S3. Randomly generated models with data-derived means and standard deviations of parameters.** (**a**) FC correlations of the randomly generated models at selected thresholds simulated at different temperatures. For each binarization threshold, we find the mean and standard deviation of the pooled group binarized time series data and sample parameters from a normal distribution with the same mean and standard deviation. (**b**) Flip rates of the same models.

Each line represents the median flip rate over all nodes, and the shaded area around it shows the range. The shaded areas are too small to be visible in most cases due to the high uniformity of the flip rates. (**c**) Progression of FC correlation for the first 32,000 steps of fitting of group models initialized with mean-binarized data-derived (blue) or randomized (orange) initial guess models. The line indicates the median, and the shaded area indicates the range. (**d**) Corresponding plot for binarization threshold 1 SD above the mean.

### S4. Individual model fitting starting from an individual data-based initial guess

As an alternative to using the group model as the initial guess for the individual models, we tested initial guesses derived from the individual data using the same method as with the group model. That is, we set $h_i = \langle\sigma_i\rangle_{\text{data}}$, $J_{ij} = \langle\sigma_i\sigma_j\rangle_{\text{data}}$, where $\langle\sigma_i\rangle_{\text{data}}$ is the mean and $\langle\sigma_i\sigma_j\rangle_{\text{data}}$ the uncentered covariance of the binarized data. We created 5 replica models for each of the 167 (testing set) subjects omitted from the group model. We then performed temperature optimization separately for each fitting target. We found that, for all thresholds, the optimal value of $\beta$ was less than 0.025, so, when fitting individual models, we searched over the range [0, 0.025], sampling 5 evenly spaced values per iteration. We then fitted the models by Boltzmann learning for 10,000 parameter updates. We compared these individual models to individual models of the same testing set subjects fitted with the group model as initial guess. For threshold 0, individualization from the group model resulted in faster convergence and essentially equivalent goodness of fit compared to starting from individual data-derived initial guesses (Figure S4a). For threshold 1, individualizing from the group model produced not only faster convergence but also better worst-case fits (Figure S4b).

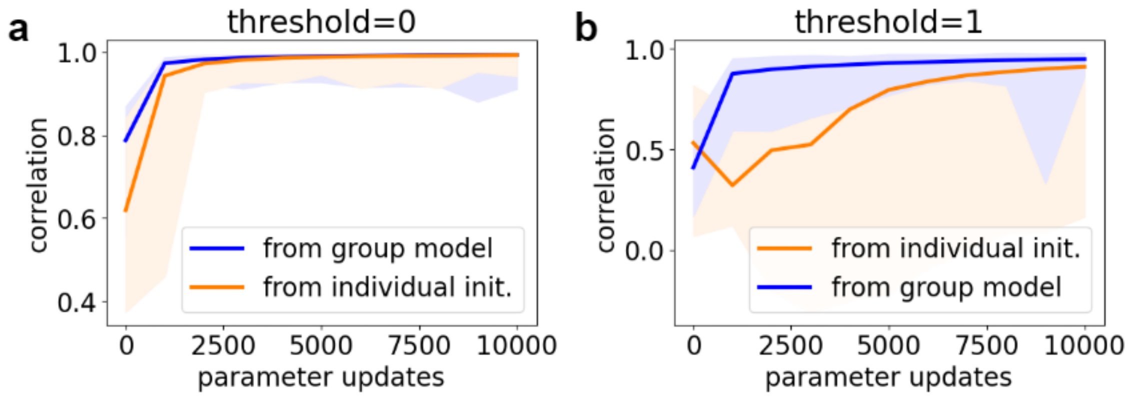


**Figure S4. Initialization of individual models with means and uncentered covariances of group models.** (**a**) Progression of fitting the Ising model to individual data binarized at threshold=0 starting from either an initial model derived from individual data or the group model. Lines represent medians, and shaded areas represent ranges over 167 individuals separate from the 670 pooled in the group data. (**b**) Corresponding figure for threshold=1.

### S5. Flip acceptance rate as a proxy for FC correlation of the initial guess model

At all binarization thresholds that allowed the initial guess group-level model to reproduce an FC close to that of the pooled group fMRI data, the value of $\beta$ that maximized FC correlation also induced a mean probability of flipping a given node state on a given step (flip acceptance rate) close to 0.5. In Supplementary Figure S5, we illustrate this with a series of plots for group models using selected binarization thresholds. Note also that the lower binarization thresholds lead to wider ranges of flip rate across regions, apparently due to the lower sensitivity of the model to individual node behavior.

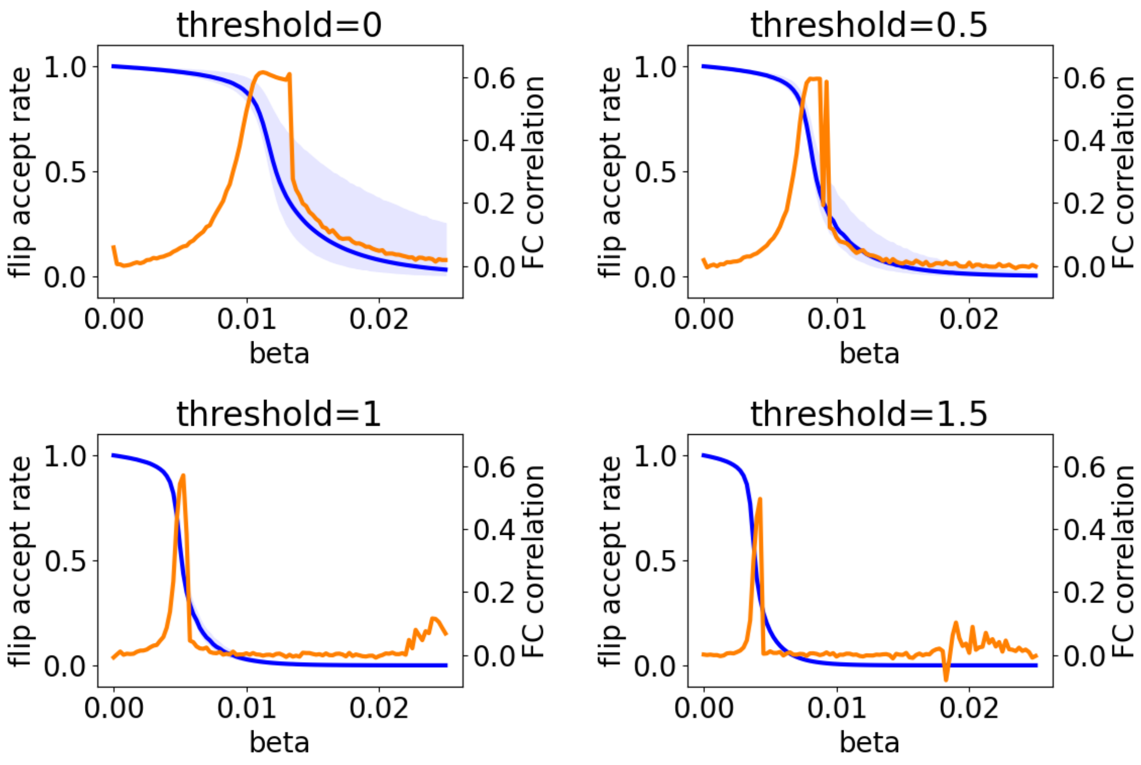


**Figure S5. Comparison of initial guess group model FC correlation to flip rate.** Each plot shows the FC correlation (orange line), mean flip rate (blue line), and range of flip rates (blue shading) of a 120,000-step Metropolis simulation of the initial guess group model (mean states for external fields and uncentered covariances for couplings) at 101 different values of $\beta$ evenly spaced between $1.0 \cdot 10^{-9}$ and 0.025 for a different binarization threshold (0.0, 0.5, 1.0, or 1.5).

### S6. Convergence of Boltzmann learning

To check how completely Ising model fitting had converged at our chosen duration of 40,000 parameter updates, we continued the fitting of each model for a total of 100,000 parameter updates. While FC correlation did increase substantially for group models at more extreme binarization thresholds, such as 1.6 SD above the mean (Figure S6a, red line), group models for data binarized at the mean (blue line) showed no significant difference between 40,000 updates (1-tailed Wilcoxon signed rank test $p = 0.64$). Models for 1 SD above the mean did show a significant difference ($p = 2.9 \cdot 10^{-15}$), but nearly all of this difference was in the outliers. The medians and 95% confidence intervals (CIs) were very similar, 0.98 versus 0.99 and [0.95, 0.99] versus [0.99, 1.0], respectively. As with the group models, the individual models binarized at the mean had already converged so thoroughly at 40,000 updates that further fitting made effectively no difference (not shown). For individual models at threshold 1SD, the FC continued to vary significantly ($p = 1.5 \cdot 10^{-89}$ for a comparison of the 40,000-update and 100,000-update models), but nearly all of that variability was in the worst fits (See Figure S6b, pale green shaded region). From 40,000 to 100,000 updates, the median increased from 0.98 to 0.99 (green line), and the 95% CI increased from [0.94, 0.99] to [0.95, 0.99] (moderately pale green shaded area).

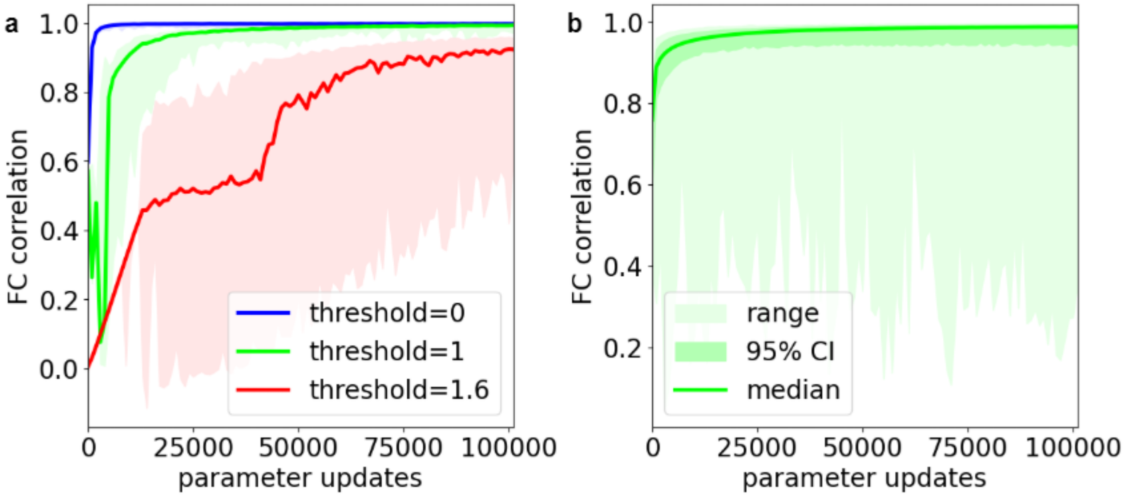


**Figure S6. Extended fitting progress plots.** (**a**) Progression of median and range of FC correlation of group models during Boltzmann learning parameter updates for the group models at 3 choices of threshold. We use the same conventions here as in Figure 3b in the main text but extend the model fitting to 100,000 parameter update steps. (**b**) Progression of median (pure green), 95% confidence interval (moderately pale green), and range (pale green) of the FC correlations of all individual models for threshold 1 SD above the mean over 100,000 parameter update steps.

### S7. Consistency of group model parameters across replica fittings

Because Boltzmann learning relies on the stochastic Metropolis algorithm (Methods subsection 2.2), model fitting runs with the same initial conditions, data-derived fitting targets, and hyperparameters ($\beta$, simulation length, and learning rate) may produce different results. To test the reliability of the Ising model fitting process, for each binarization threshold that we applied to the 670-subject group data, we compared not only FC correlation (Figure 3c in main text) but also parameter variability across the 101 replica fittings. In Figure S7a, we selected the replica that produces the best correlation between simulated and data FC and then computed the correlation between the parameters of that best model and those of each other replica. At thresholds below 1, correlations for both $h_i$ and $J_{ij}$ (elements above the diagonal) vectors were always over 0.99. At 1, all $J_{ij}$ correlations remained at least 0.99, and most $h_i$ correlations remained above 0.99, but the first outlier with a correlation of only 0.82 appeared. Even at much higher thresholds, the lowest replica model $J_{ij}$ correlation observed was just under 0.98. While the minimum $h_i$ dropped much lower in several cases, the median correlation remained above 0.98 until the threshold reached 2.5. This shows that, overall, the structure of the fitted group model parameter vectors retained a high degree of consistency across independent replica fittings.

However, replica fittings did still show some variability, as shown in Figure S7b. Here, we first took the SD over all replica values of a given single region $h_i$ or single region pair $J_{ij}$ and then plotted how the median and range of SD over regions or region pairs depended on threshold. Because we chose a different $\beta$ value at each threshold prior to fitting, we multiplied $\beta$ into both $h_i$ and $J_{ij}$ prior to analyzing the SD of the parameters. This placed the parameters of all models on the same scale in terms of influence on model behavior in the Metropolis simulation. For thresholds up through 0.8, all SD values over replicas remained smaller than 0.001. For reference, all best replica models in this range had both $h_i$ and $J_{ij}$ parameters with means on the order of 0.001 and SD over regions or region pairs on the order of 0.01. At threshold 1, most SD over replicas remained on the order of 0.001 or smaller. At thresholds above 1, some thresholds developed larger SD, but these increases occurred sporadically. In these cases, even as the median SD increased, the range remained small, and SD of $h_i$ and $J_{ij}$ increased together.

Figure S7c illustrates why this is the case. The overall pattern of differences among $h_i$ elements remains the same for all replicas at threshold 1.6, but three outliers have drifted downward relative to the majority. Figure S7d shows that the behavior of the model was robust to this drift effect, as the FC correlations (compared to the data FC) of the outlier models were near the middle of the distribution. The corresponding figures for $J_{ij}$ are visually similar but with less drift. This is unsurprising when one considers that the behavior of the Metropolis simulation depends on the differences between entropies of states rather than their absolute entropies (See Methods). Specifically, in Equation 1, we have $s(\vec{\sigma}) = \sum_{i=1}^{N}(h_i\sigma_i + \sum_{j}^{N} J_{ij}\sigma_i\sigma_j)$. At high thresholds, at most time points, $\sigma_j = -1$ for most $j$. Consequently, $s(\vec{\sigma}) \approx \sum_{i=1}^{N}\sigma_i(h_i - \sum_{j}^{N} J_{ij})$. If all $J_{ij}$ drift by some amount $d$, and all $h_i$ drift by some larger amount $c = Nd$, then this sum remains the same.

Figure S7e and S7f show how SD over replicas changed over the course of fitting for $h_i$ and $J_{ij}$, respectively. At threshold 0 (blue), all SD drifted upward slightly at first but quickly leveled off and remain small. At 1 (green), SD of $h_i$ and $J_{ij}$ both increased early in fitting but stop and began to decrease at different points. At 1.4 (orange), both SD quickly increased at an arbitrary point in fitting and then decreased again. At 1.6 (red), all SD of $h_i$ grew large but eventually leveled off, but SD of $J_{ij}$ had some outliers that continued to grow. Overall, this shows that, at small thresholds, the fitting was highly repeatable with almost no variability across replicas. At thresholds around 1, random drift affected the parameters, but they still eventually converged to consistent values. At higher thresholds, fitting was more chaotic but the largest differences in replicas were due to uniform drift of parameters instead of independent parameter changes, meaning that most replicas still remain highly correlated with each other and contain the same information.

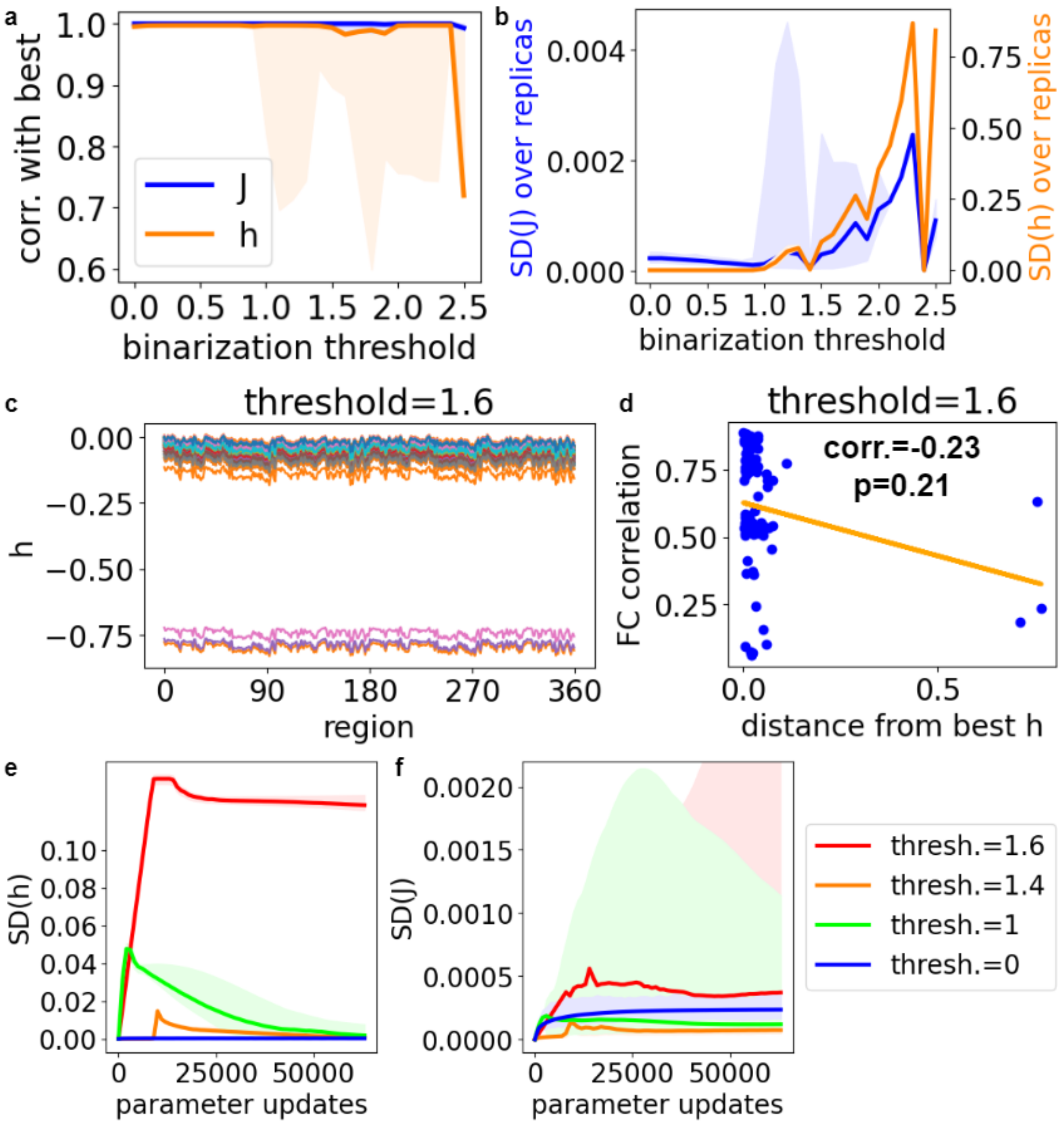

**Figure S7. Parameter variation across replica fittings of the group Ising model.** (**a**) Correlation between a given replica $h_i$ (orange) and $J_{ij}$ (blue, upper triangular part) parameter vector and that of the replica that gives the highest correlation between data and model FC, taken over 101 independent fittings to the same 670-subject group data binarized at different thresholds. The line shows the median, and the shaded area shows the range over. (**b**) SD of $h_i$ (orange) or $J_{ij}$ (blue) taken over the 101 replicas. The line shows the median, and the shaded area shows the range over all 360 regions for $h_i$ or all 64,620 region pairs for $J_{ij}$. (**c**) Illustration of drift in $h_i$ of group models. Each line represents the $h_i$ vector of a single replica. (**d**) Scatter plot of distances between individual replica $h_i$ vectors and correlation between data and model FC. Each point represents a replica fitted to group data at threshold 1.6. (**e**) Change in SD of $h_i$ over the course of fitting to group data binarized at selected thresholds. The line shows the median, and the shaded area shows the range over all 360 regions for $h_i$. (**f**) Change in SD of $J_{ij}$ during fitting. The line shows the median, and the shaded area shows the range over all 64,620 region pairs.

### S8. Comparison of group model parameters between thresholds

To test to what extent the Ising models retained a common structure at different thresholds, we computed the correlation between parameters in models fitted to data binarized at threshold 1 and all other thresholds (0 through 1.5) (Figure S8a). While FC correlation drops with increasing threshold over this range (See Figure 3c in the main text.), the median FC correlation at threshold 1.5 is still 0.822, indicating a good enough fit for this comparison. To avoid artificially inflating the correlations with multiple nearly identical data points, we took the mean over replicas of each model before taking the correlations. A noted above, we multiply $\beta$ into both $h_i$ and $J_{ij}$ to place the model parameters on the same effective scale for all thresholds. For $J_{ij}$, the correlations with the threshold 1 model remain above 0.75 across all thresholds. For $h_i$, the correlations with the threshold 1 model remain above 0.75 across nearly all thresholds, the exception being 0. This shows that the pattern of connectivity of the group model is stable across a wide range of thresholds. It also shows that choosing a non-0 threshold qualitatively changes the pattern of $h_i$ values but that the pattern that non-0 thresholds induce remains stable over nearly the same range of thresholds as $J_{ij}$. However, we see that changing the threshold changes the overall scales of $h_i$ and $J_{ij}$, as measured by their means, and the overall scales of their respective heterogeneities, as measured by their SDs (Figure S8b). Whereas the mean of $J_{ij}$ remains nearly the same as its SD gradually shrinks (blue), the mean of $h_i$ starts near 0 and decreases, slowly and steadily at first and then faster and more erratically as the binarization threshold increases past 1. The SD of $h_i$ grows to a maximum close to a threshold of 0.5 before gradually shrinking again. At our chosen thresholds, 0 and 1, that we subsequently use for individual models, the group model values of $h_i$ are completely uncorrelated between the two thresholds (Figure S8c), while the values of $J_{ij}$ pairs are highly correlated (Figure S8d). From the scales of the plots, we can see that increasing the threshold from 0 to 1 induces a dramatic increase in the heterogeneity of $h_i$ while leaving $J_{ij}$ largely the same.

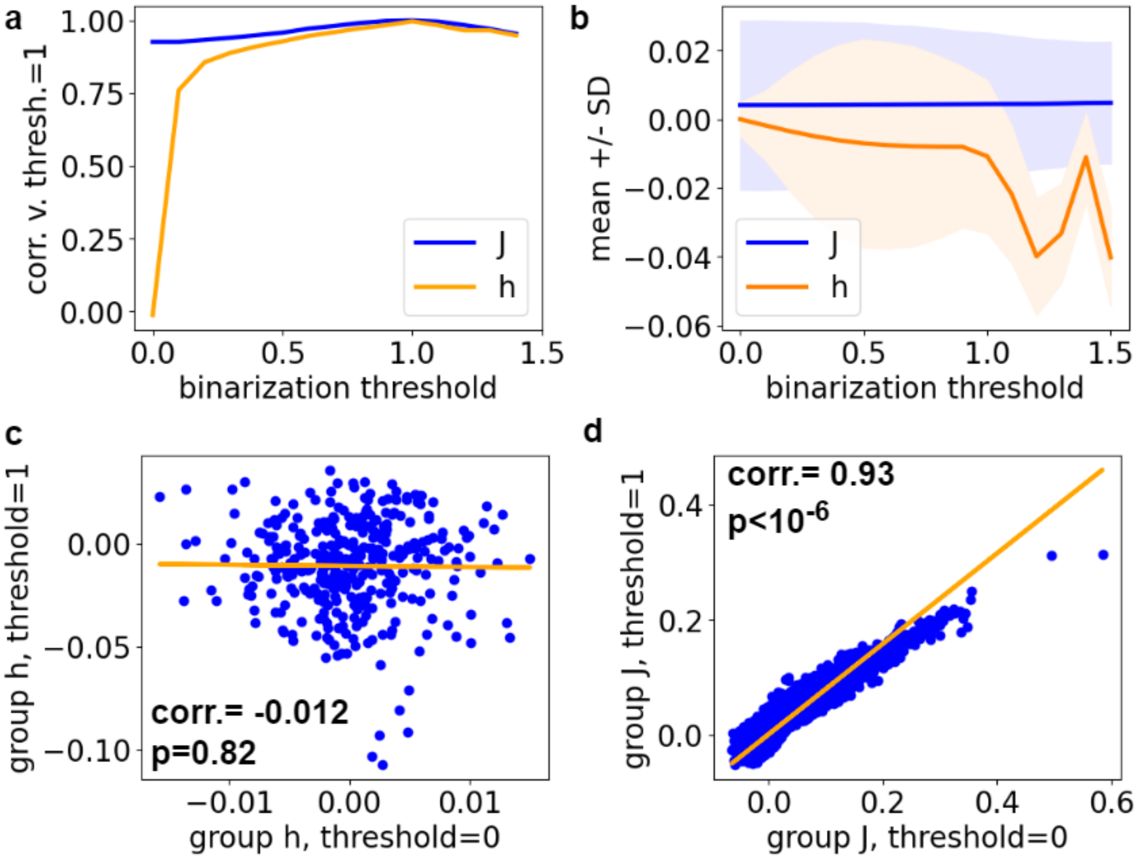


**Figure S8. Correlations between corresponding group model parameters at different thresholds.** (**a**) Correlation between $h_i$ (orange) or $J_{ij}$ (blue) at threshold 1 and at the given threshold (x-axis). (**b**) Mean (line) and SD (shaded area) of parameters $h_i$ over regions (orange) or $J_{ij}$ over region pairs (blue). (**c**) Scatter plot (blue) and least squares linear regression (orange) comparing group model $h_i$ at thresholds 0 and 1. Each data point represents a single region. (**d**) Scatter plot (blue) and least squares linear regression (orange) comparing group model $J_{ij}$ at thresholds 0 and 1. Each data point represents a single pair of regions.

### S9. Comparison of individual model parameters between thresholds

We next considered the ensembles of 837 models fitted to individual subject data binarized at either 0 or 1. We first compared the local mean over all subjects at each region or region pair. As noted with the group model values, the local mean values of $h_i$ had no correlation across thresholds (Figure S9a), while the local means of $J_{ij}$ were highly correlated (Figure S9b). The SDs of $h_i$ at the two thresholds moderately but significantly correlated with each other (Figure S9c). This showed that the models were qualitatively different at the level of individual variation as well as at the level of region-to-region variation but that some of the same regions had relatively low individual variation at either threshold. By contrast, the SDs of $J_{ij}$ were highly correlated (Figure S9d), indicating that the region pairs with high individual variatiability of $J_{ij}$ remained the same at either threshold.

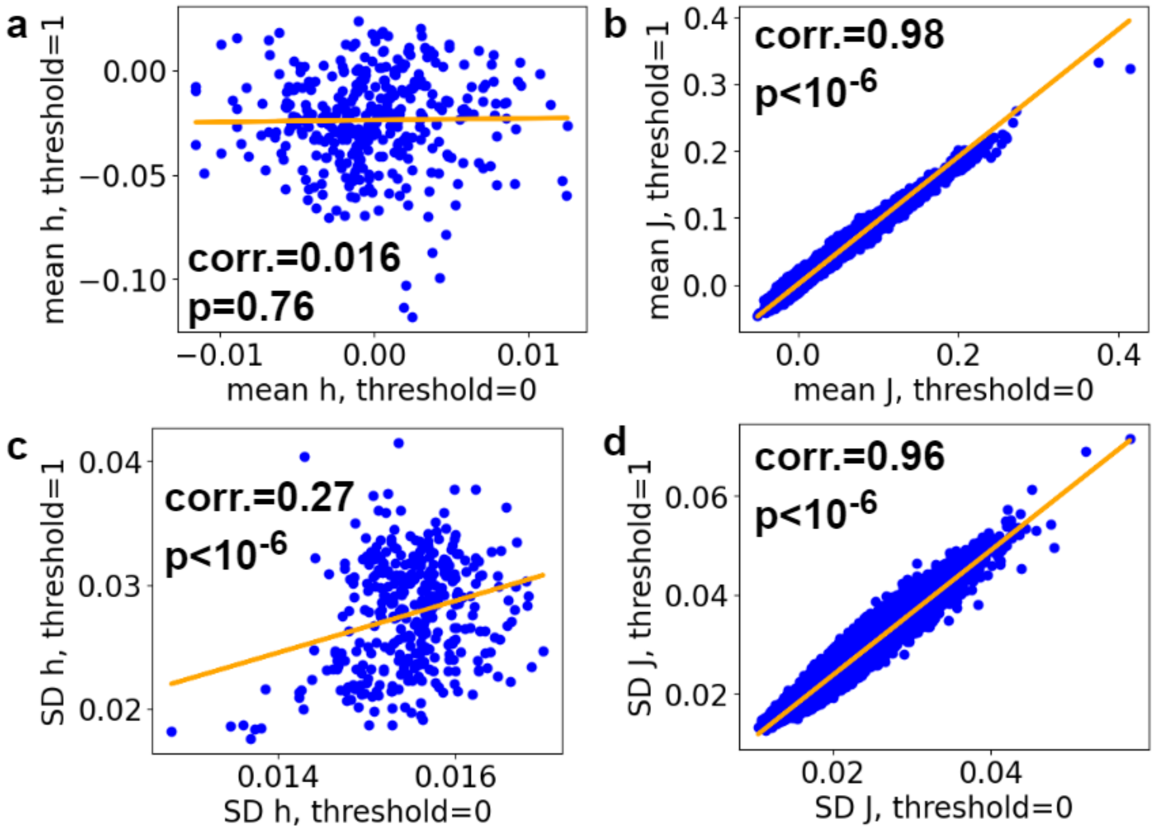


**Figure S9. Correlations between corresponding individual model parameters at different thresholds.** (**a**) Comparison of corresponding mean $h_i$ values over all subjects for thresholds 0 and 1. Each point represents a region. (**b**) Comparison of corresponding mean $J_{ij}$ values over all subjects for thresholds 0 and 1. Each point represents a region pair. (**c**) Comparison of the SD of $h_i$ over all subjects for thresholds 0 and 1. Each point represents a region. (**d**) Comparison of the SD of $J_{ij}$ values over all subjects for thresholds 0 and 1. Each point represents a region pair.

### S10. Cross-validation of group-level structure-to-parameter least-squares regression models

To test the robustness of the region-to-region correlations between the group mean structural features and group Ising model parameters, we cross-validated the least-squares linear regressions. For each correlation, we performed 10,000 even train-test splits. For the linear model predicting $J_{ij}$ from SC, we randomly split the region pairs into 32,310 training pairs and 32,310 testing pairs (Figure S10a). For comparison, we repeated this process with prediction of binarized group data FC from SC (Figure S10b). For the linear model predicting group model $h_i$ from thickness, myelination, curvature, and sulcus depth, this meant splitting the brain into 180 training set regions and 180 testing set regions (Figure S10c). For comparison, we repeated this process with prediction of binarized group data mean state (Figure S10d).

Figure S10a shows that the distributions of training (blue) and testing (orange) $J_{ij}$ prediction correlations were nearly identical at all binarization thresholds (separate Wilcoxon signed rank tests, all $p > 0.05$) and that the variability of the correlations over different samplings of region pairs at the same threshold was generally small (all SD < 0.01 for both training and testing). Likewise, the distributions of training and testing FC prediction correlations were nearly identical (all $p > 0.05$) and all had SD < 0.01. This shows that the population-level relationships between brain-wide patterns of $J_{ij}$, FC, and SC are robust and generalize well when estimated from one subset of region pairs and then used to predict $J_{ij}$ or FC from SC for others.

The models predicting $h_i$ from structural features generalized less well from training to testing region sets in that testing prediction correlations were consistently lower (separate 1-sided Wilcoxon signed rank test for each threshold, all $p < 10^{-100}$) (Figure S10c). However, the means of the distributions of training and testing correlations followed the same upward trend with increasing binarization threshold, and the largest difference between training and testing mean correlations is 0.11. In general, the means approach each other as the threshold increases, with the difference shrinking to 0.052 at threshold 1. At each threshold, SD (shaded area) is similar over both training and testing regions, while mean correlation (center line) is lower for testing than for training. Differences between training and testing means vary from 0.025 to 0.11 with the difference at threshold 1 being 0.052. The prediction correlation for mean state showed a similar downward shift from training to testing while still following the same pattern of change versus the binarization threshold (Figure S10d). Overall, these results show that, while the use of multiple structural features may lead to some mild overfitting, the general population-level relationships between $h_i$ and structural features are robust, especially at thresholds where the permutation tests showed them to be significant (See Figure 4b in main text).

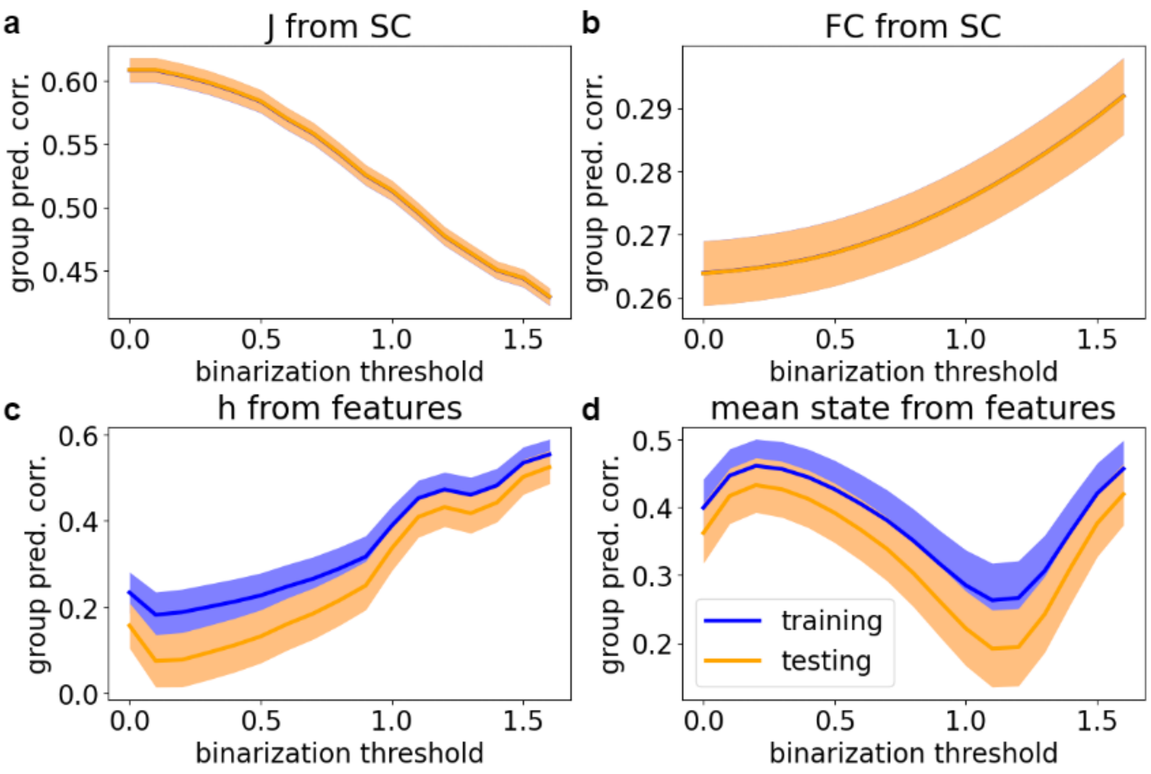


**Supplementary Figure S10. Comparison between training and testing prediction correlations from group-level least squares regression models.** In each figure, the vertical axis represents the correlation between the group Ising model parameter ($h_i$ or $J_{ij}$) and the parameter value predicted from structural features. The blue line and shaded area represent the mean and SD of the training correlation, and the orange line and shaded area represent the mean and SD of the testing correlation. We took the mean and SD over 10,000 random 50-50 train-test splits of the regions (for $h_i$) or region pairs (for $J_{ij}$) at each binarization threshold. (**a**) Correlation between the Ising model $J_{ij}$ and $J_{ij}$ predicted from SC. Training (blue) and testing (orange) bands completely overlap. (**b**) Correlation between the FC of the binarized data and FC predicted from SC. Training (blue) and testing (orange) bands completely overlap. (**c**) Correlation between the Ising model $h_i$ and $h_i$ predicted from the four structural features. (**d**) Correlation between the mean state of the binarized data and mean state predicted from the four structural features.

### S11. Extending brain-wide trends in structure-function relationships from group data to individual data

To test the extent to which the whole-brain level relationships between group model $h_i$ and group mean structural MRI features or between group model $J_{ij}$ and group mean SC applied at the level of individual subjects, we used the linear models that we previously fitted to group Ising models and group mean structural data and used them to predict Ising model parameter values from individual structural feature data. We then calculated one prediction correlation per subject over all 360 regions or 64,620 region pairs. We compared distributions of correlations for the individual models at both thresholds used for individual models in the main results, 0 and 1, and compared results for $h_i$ to results for binarized data mean state and compared results for $J_{ij}$ to results for binarized data FC.

Figure S11 a and b show that, at both binarization thresholds, individual-level prediction correlations of $J_{ij}$ from SC were substantially worse than the group-level correlation but still universally better than predictions of FC from SC. Figure S11 c and d show that, whereas the group-level prediction correlations of $h_i$ at threshold 1 and mean state at threshold 0 from all four structural MRI features were similar, the $h_i$ prediction model generalizes significantly better to individual data (separate 1-tailed Wilcoxon signed rank tests for training and testing subjects, both $p < 10^{-6}$). As in all other results, prediction correlations for $h_i$ at threshold 1 were greater than at threshold 0 (both 1-tailed Wilcoxon signed rank test $p < 10^{-6}$). In all eight train-test splits, the distributions of correlations for the 670 training subjects originally included in the group mean data were not significantly larger than those for the 167 testing subjects (1-tail Mann-Whitney U-tests, smallest $p = 0.063$). Overall, these results suggest that, while the group-level whole-brain correlations between structure and function do not fully translate to individual data, the correlations found for Ising model parameters translate from group to individual better than do those for simple statistics computed directly from the binarized fMRI data.

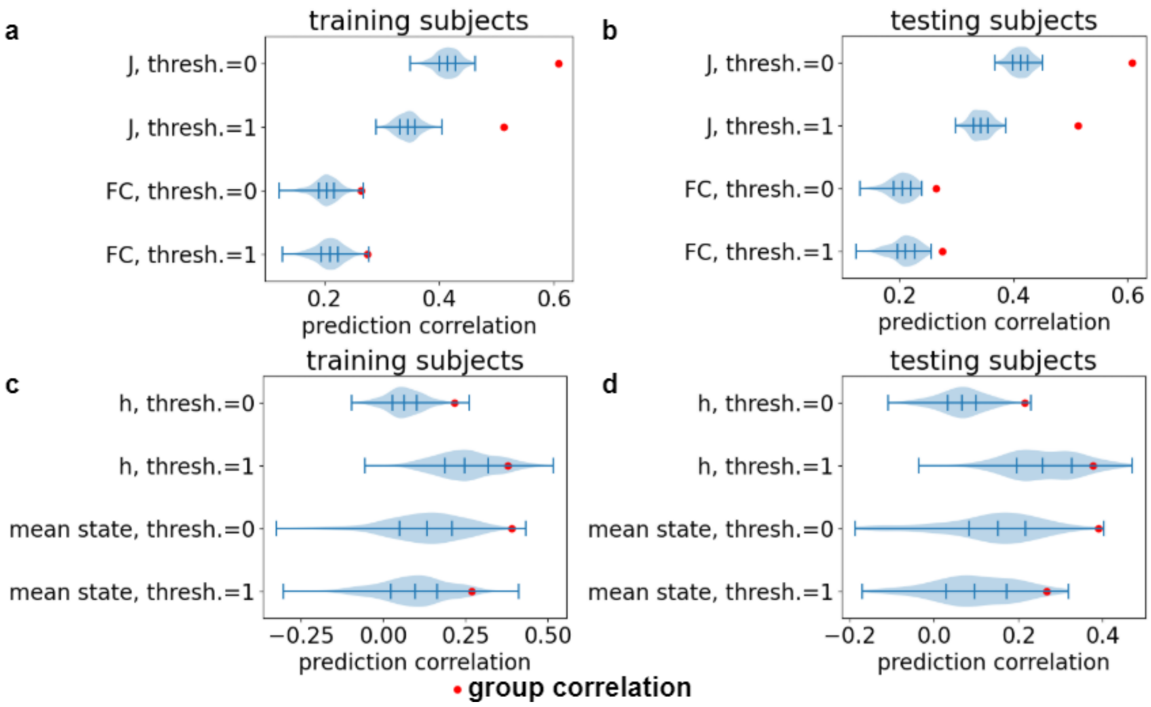


**Supplementary Figure S11. Applying structure-to-parameter linear models from group mean structural features and group Ising models to individual data and Ising models**. Correlations between individual Ising model parameters and parameters predicted from individual structural feature data using the least squares regression models fitted to group-level Ising models and group mean structural data. The red dot indicates the group model prediction correlations. (**a**) Prediction of $J_{ij}$ or FC from SC applied to training data subjects. (**b**) Prediction of $J_{ij}$ or binarized data FC from SC applied to testing data subjects. (**c**) Prediction of $h_i$ or binarized data mean state from all four structural features applied to training data subjects. (**d**) Prediction of $h_i$ or binarized data mean state from all four structural features applied to testing data subjects.

### S12. Direct correlations between structural features and Ising model parameters

To test how strongly individual structural features corresponded to $h_i$ and to illustrate the value of combining information from multiple structural features, we compared the multiple least squares prediction correlations to direct correlations between model parameters and structural features. Figure S12a shows how the correlation between the group mean of each region feature and group model $h_i$ changes with binarization threshold. For comparison, Figure S12b shows the direct correlation between each group mean region feature and the mean state calculated from the binarized group data. While the maximum absolute values of the mean state and $h_i$ correlations are similar, the $h_i$-feature correlations follow more nearly monotonic upward or downward trends, each eventually achieving significance. By contrast, the mean state correlations not only change direction as the threshold increases but reach significance in one direction, then the other. This suggests that $h_i$ has a simpler, more interpretable relationship with local region excitability than does the mean state.

Figures S12 c-f show the distributions of direct correlations between individual model $h_i$ or individual data mean state and individual structural features. In all plots, $h_i$ at threshold 1 has the largest correlations. Whereas the group mean state four-feature prediction correlation is greatest at threshold 0.2, the median individual prediction correlation is greatest at threshold 0. We have not included plots of the distributions at all thresholds, because they are visually similar.

The subsequent plots show the distributions of direct correlations between local individual differences in model parameters and individual features. Figure S12g compares direct correlations between SC and $J_{ij}$ or SC and binarized data FC at threshold 0 or 1. All four distributions are roughly similar, though $J_{ij}$-SC correlation has larger outliers than FC-SC correlation at either threshold. In all four cases, only positive correlations achieve statistical significance, indicating that the negative correlations are spurious.

Whereas myelination has the strongest and curvature the weakest group-level correlation at threshold 1 (Figure S12a), different individual difference correlations are strongest at different regions (Figure S12h). While myelination has the largest outlier, curvature has the largest median and is the most common strongest correlation. Both curvature and sulcus depth tend to have stronger correlations than thickness and myelination (1-tailed Wilcoxon signed-rank test of absolute values, all four $p < 0.001$). Curvature also tends to have slightly stronger correlations than sulcus depth, though the difference falls just short of significance ($p = 0.0013$). This suggests that myelination and cortical folding play different roles in differentiating the dynamics of brain regions, with myelination having the most importance to the whole-brain functional gradients common to all subjects and cortical folding having more relevance to local individual differences that develop within the constraints that myelination sets.

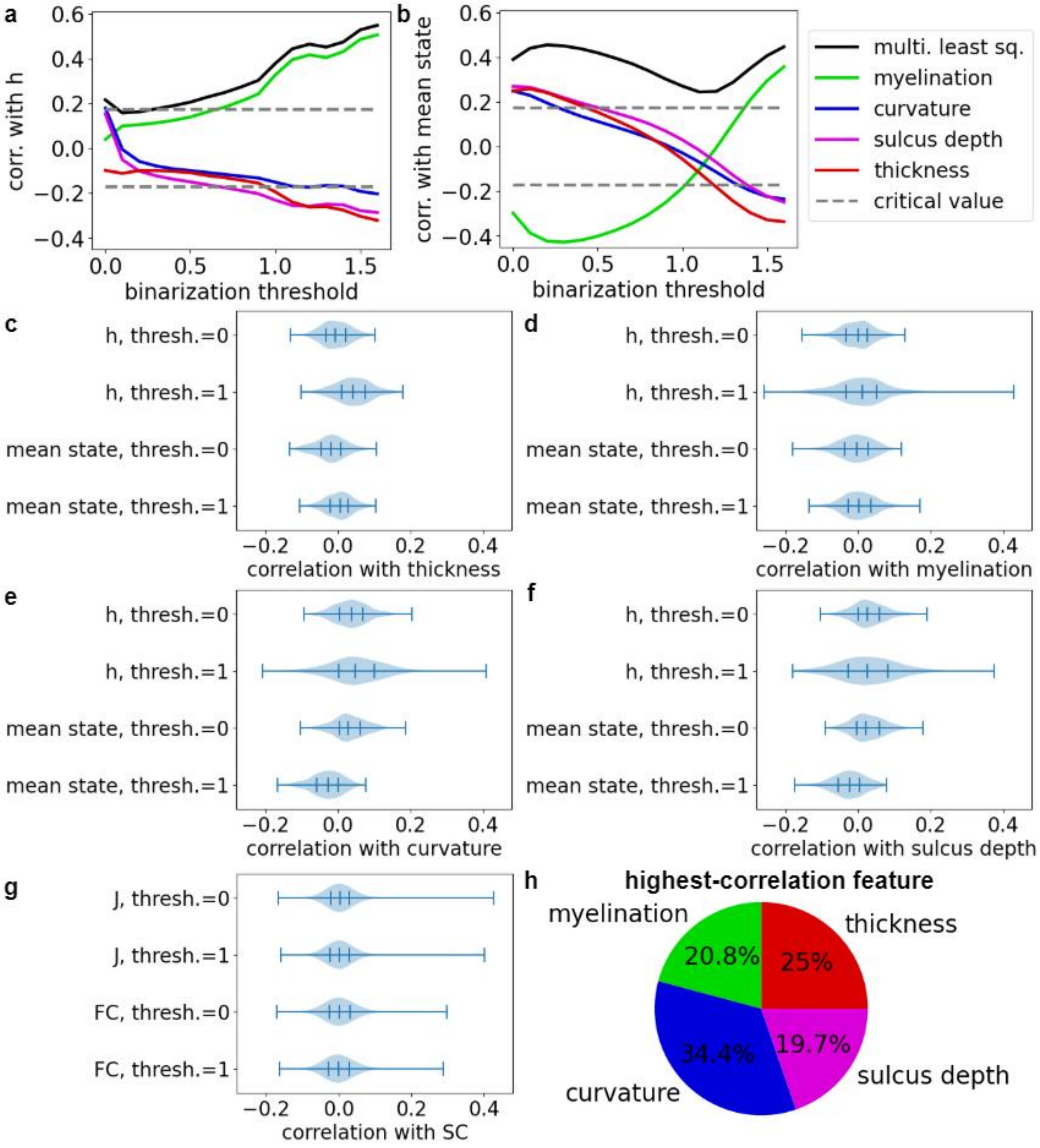

**Supplementary Figure S12. Direct correlations between Ising model parameters and structural features.** (**a**) Direct correlations between structural features and group model $h_i$ as a function of binarization threshold. The black curve labeled "multi. least sq." represents the correlation between $h_i$ and the multiple least squares regression predicting $h_i$ from all four features. Because correlations can be positive or negative, we mirror the largest critical value over the horizontal axis, indicating that values that fall outside the paired broken lines are significant. (**b**) Direct correlations between structural features and mean state computed from the binarized group data as a function of binarization threshold. The black curve labeled "multi. least sq." represents the correlation between the mean state and the multiple least squares regression predicting the mean state from all four features. Since the group model $J_{ij}$-SC and group data FC-SC correlations are always positive, direct correlations and least squares regression model correlations are identical. (**c**)-(**f**) See Figure 4a. Violin plots of direct correlations between $h_i$ or mean state at threshold 0 or 1 and a single feature: (**c**) thickness (**d**) myelination. (**e**) curvature (**f**) sulcus depth. Each individual data point is the correlation over all individuals for a region. Tick marks represent quartiles. (**g**) Violin plots of direct correlations between individual differences in SC and Ising model $J_{ij}$ or individual binarized data FC at threshold 0 or 1. Each individual data point is the correlation over all individuals for a connection between a pair of regions. Tick marks represent quartiles. (**h**) Pie plot indicating the percentage of regions for which each feature has the highest-absolute-value correlation with $h_i$.

### S13. Cross-validation of individual difference structure-to-parameter least-squares regression models

To test how consistent relationships between model parameters and structural features are across subjects, we performed cross-validations of the individual difference least squares regression linear models. We used 100 randomized splits of each set of individual Ising models into 670 training subjects and 167 testing subjects, approximating an 80-20 split. We fitted local individual-level linear models to the Ising model parameters and structural features of the training subjects as described in the main text and then applied those models to the testing subjects. For each region or pair of regions, this resulted in a sampling of 100 pairs of training and testing prediction correlations.

Figure S13 shows that the means of training and testing prediction correlations (blue lines) closely align, but the SD (orange error bars) values indicate substantial variation across different splits of the data. Correlations between $J_{ij}$ or FC and their values predicted from SC tended to have especially large variability but only in cases where the mean correlations were weak. These represent cases where SC and values predicted from it had very little variance, making the prediction correlations numerically unstable. See Figure 5a in the main text.

Of particular interest is whether the regions or region pairs that had significant correlations according to the permutation tests reported in the main text have structure-parameter relationships that generalize to new subjects. For the 96 regions that had significant prediction correlations for $h_i$ predicted from all four features, the median difference between training and testing prediction correlations was 0.022, compared to a median training subject correlation of 0.20 and median testing subject correlation of 0.19. The overall fraction of permutations in which training correlation exceeded testing correlation was 0.60, and a 1-tailed Wilcoxon signed rank test indicated that testing correlations were significantly lower ($p < 10^{-6}$). Overall, this indicates the presence of significant but moderate overfitting and suggests that, for these regions with relatively strong structure-function coupling, individual differences in structure have a consistent direction of influence on local excitability, here represented by $h_i$.

By contrast, when we selected the 495 region pairs for which the correlations between $J_{ij}$ of Ising models for threshold 1 and $J_{ij}$ predicted from SC were significant, we found that deviations between training and testing correlations were roughly symmetric around 0 (2-tailed Wilcoxon signed rank test $p = 0.081$). The median difference between training and testing correlations was 0.00029, compared to a median training correlation of 0.17 and median testing correlation of 0.18. This indicates that, at least for this small fraction of connections with strong structure-function coupling, the effect of individual differences in SC on individual differences in $J_{ij}$ generalizes well across different subsets of subjects. One common feature of all plots in Figure S12 is that, as both training and testing correlations increase, their means approach the identity line (shown in black), indicating that the disparity between them vanishes. This implies that the strongest structure-function relationships are also the ones that hold true most consistently across different subsets of individuals.

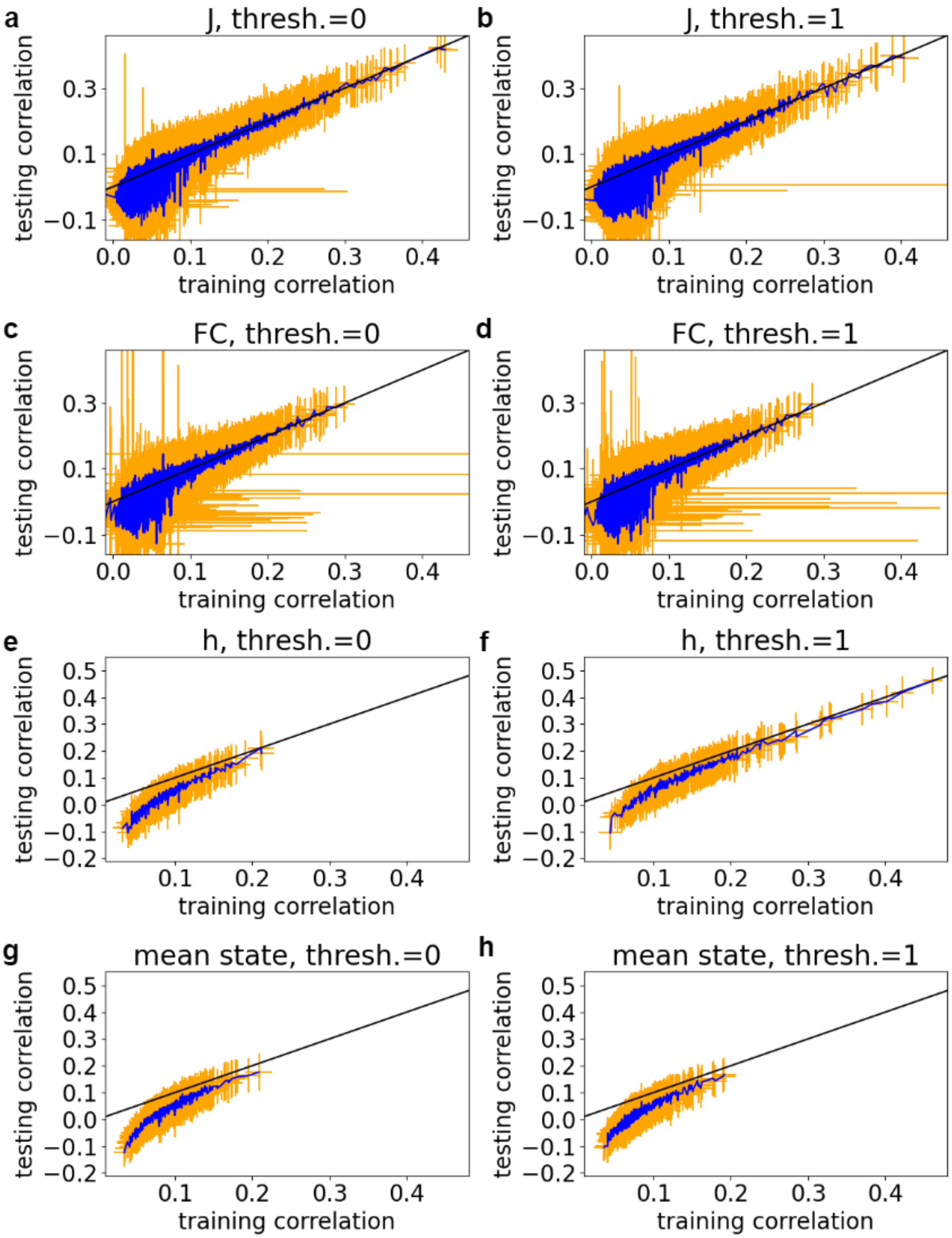


**Supplementary Figure S13. Comparison between training and testing prediction correlations from the least squares regression models.** The blue line represents the means, and the orange error bars represent the SD over 100 random train-test splits. The black line is the identity line, plotted for reference. (**a**) Correlation between Ising model $J_{ij}$ at threshold 0 and $J_{ij}$

predicted from SC. (**b**) Correlation between Ising model $J_{ij}$ at threshold 1 and $J_{ij}$ predicted from SC. (**c**) Correlation between the FC of the data binarized at threshold 0 and the FC predicted from SC. (**d**) Correlation between the FC of the data binarized at threshold 1 and the FC predicted from SC. (**e**) Correlation between Ising model $h_i$ at threshold 0 and $h_i$ predicted from the four structural features. (**f**) Correlation between Ising model $h_i$ at threshold 1 and $h_i$ predicted from the four structural features. (**g**) Correlation between the mean state of the data binarized at threshold 0 and the mean state from the four structural features. (**h**) Correlation between the mean state of the data binarized at threshold 1 and the mean state from the four structural features.

### S14. Correlations between Ising model external field and NODDI, DTI features

To compare the T1/T2 ratio to other myelin-associated signals, we compared group-level region-wise trends and local individual differences in $h_i$ to DTI features derived using the Accelerated Microstructure Imaging via Convex Optimization (AMICO) pipeline (Daducci, 2015) and Neurite Orientation Dispersion and Density Index (NODDI) model (Zhang, 2012). See (Tromp, 2023) for a brief explanation of DTI features not specific to NODDI: fractional anisotropy (FA), radial diffusivity (RD), axial diffusivity (AD), and mean diffusivity (MD). As shown in (Jespersen, 2010) and referenced in (Zhang, 2012), neurite density correlates with myelination, so we use the neurite density index (NDI) measure from NODDI. By contrast, increased isotropic volume fraction (ISOVF), an estimate of the fraction of the volume occupied by cerebrospinal fluid (Zhang, 2012), implies a lower fraction of volume occupied by neurites and myelin.

Figure S14a shows how correlations between group model $h_i$ and these features changed with fMRI binarization threshold. We plot the structural MRI features used in the main text alongside them for comparison. While none of the correlations with non-NODDI DTI features correlated significantly at any threshold, both NODDI features achieved significant correlations at thresholds around 1. Group mean values of RD, AD, and MD correlate strongly with each other: 0.98 for MD v. RD, 0.95 for MD v. AD, and 0.86 for RD v. AD. Consequently, all three show nearly identical correlations with group model $h_i$ across all thresholds. FA is among the weakest correlations at all thresholds. ISOVF exhibited the expected trend, starting out non-significant and showing increasingly strong negative correlations at higher thresholds. Unexpectedly, NDI also showed significant negative correlations with $h_i$, even at binarization thresholds well below those at which other features achieved significance.

Figure S14b shows the correlation between $h_i$ and predicted $h_i$ from multiple least-squares regression for different combinations of features, either the T1- and T2-ratios from structural MRI (MRI), the non-NODDI DTI features (DTI), the NODDI features (NODDI), the DTI and NODDI features together (DTI+NODDI), or all ten features together (MRI+DTI+NODDI). All prediction correlations trend upward with threshold and achieve significance within the range of thresholds considered. NODDI and non-NODDI DTI features together achieve much stronger correlations than those from NODDI features or non-NODDI DTI features alone at all thresholds and even exceed the correlations from the combined non-DTI MRI features. These combined DTI+NODDI prediction correlations are significant at all thresholds. Combining all 10 features produces even stronger correlations at all thresholds.

Figure S14c shows distributions of local individual difference correlations with individual features and multiple least-squares regression predictions for $h_i^{(s)}$ fitted to data binarized at threshold 1 (c). Unlike with the group-level global correlations, $h$(MRI) prediction correlations tend to be stronger than $h$(DTI+NODDI) prediction correlations (2-sided Wilcoxon signed-rank test $p = 5.2 \cdot 10^{-20}$). Combining all 10 features achieves a significant improvement over only DTI+NODDI features (1-sided Wilcoxon $p = 7.9 \cdot 10^{-61}$) or only MRI features (1-sided Wilcoxon $p = 4.2 \cdot 10^{-59}$). However, the number of significant MRI+DTI+NODDI correlations (50) is slightly lower than the number of significant MRI-only correlations (55), suggesting that some of the increases in correlations may be due to overfitting. Single-feature DTI and NODDI feature correlations generally fall within the same range as those for T1- and T2-weighted MRI features, with the bulk of each distribution falling within [-0.2, 0.2] and outliers ranging from -0.33 (NDI) to 0.39 (AD).

Figure S14d shows the corresponding distributions of correlations for threshold 0 Ising models. Correlations for threshold 1 are significantly greater than for threshold 0 for every feature or combination of features (maximum $p = 0.0020 < 0.0031$). The threshold 1 models also have more regions with significant correlations for every feature or combination of features. At threshold 0, most have no significant correlations, the exceptions being curvature (1 significant), sulcus depth (3), and $h$(MRI).

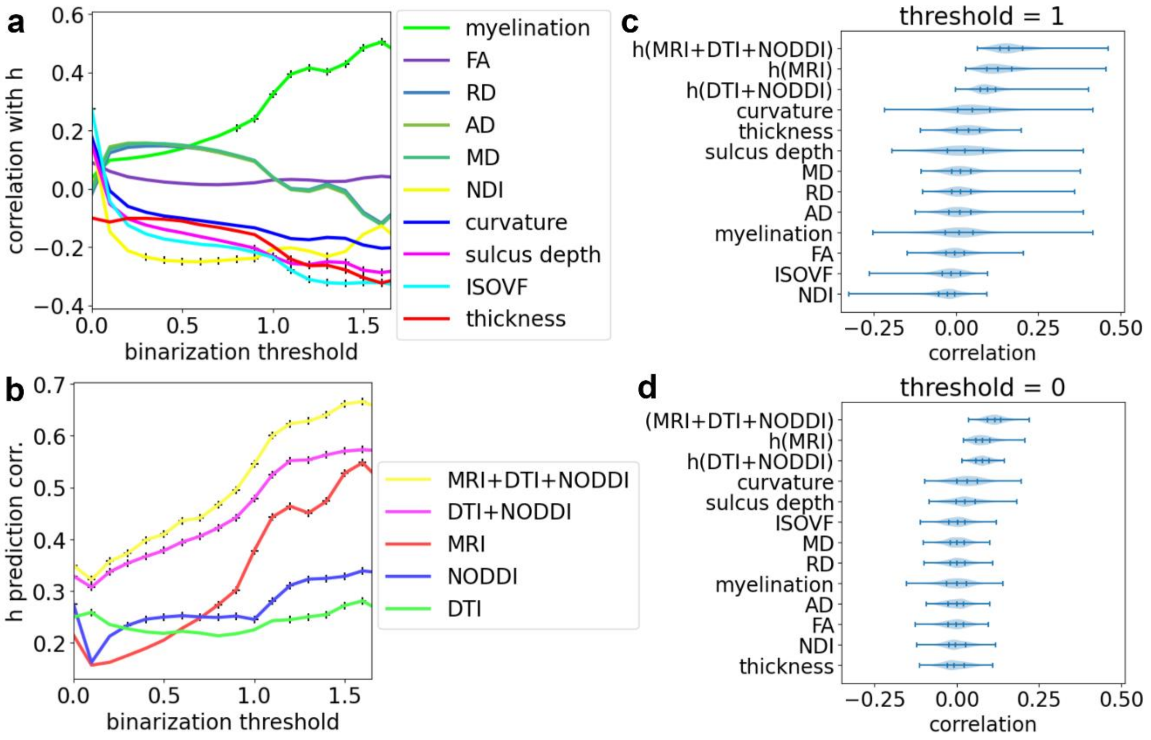


**Supplementary Figure S14. Correlations between external field parameter and NODDI, DTI features.** (**a**) Region-wise correlation between group Ising model $h_i$ and local mean structural features. Features: thickness, myelination (T1w/T2w ratio), curvature, and sulcus depth from structural MRI; fractional anisotropy (FA), radial diffusivity (RD), axial diffusivity (AD), and mean diffusivity (MD) from basic DTI processing; neurite density index (NDI) and isotropic volume fraction (ISOVF) from NODDI processing of DTI data. Black '+' markers indicate significant correlations. (**b**) Region-wise multiple least-squares regression prediction of $h_i$ using different combinations of features from T1- and T2-weighted structural MRI (thickness, myelination, curvature, sulcus depth), basic DTI processing (FA, RD, AD, MD), and NODDI DTI processing (NDI, ISOVF). Black '+' markers indicate significant correlations. (**c**) Distributions of subject-wise correlations between individual Ising model $h_i^{(s)}$ and individual structural feature values at each region for Ising models fitted to data at threshold 1. Feature abbreviations are the same as in plot a. Plots are in descending order of median (middle notch). (**d**) Corresponding plot of individual $h_i^{(s)}$-feature correlations for Ising models fitted with threshold 0.
